# Optimized dimerization of the PAR-2 RING domain drives cooperative and selective membrane recruitment for robust feedback-driven cell polarization

**DOI:** 10.1101/2023.08.10.552581

**Authors:** Tom Bland, Nisha Hirani, David Briggs, Riccardo Rossetto, KangBo Ng, Neil Q. McDonald, David Zwicker, Nathan W. Goehring

## Abstract

The behavior of cell polarity networks is defined by the quantitative features of their constituent feedback circuits, which must be tuned to enable robust and stable polarization, while also ensuring that networks remain responsive to dynamically changing cellular states and/or spatial cues that arise during development. Using the PAR polarity network as a model, we demonstrate that these features are enabled by dimerisation of the polarity protein PAR-2 via ubiquitin-independent function of its N-terminal RING domain. Specifically, we combine theory and experiment to show that dimer affinity is optimized to achieve dynamic, selective, and cooperative recruitment of PAR-2 to the plasma membrane during polarization. Reducing dimerization results in loss of positive feedback and compromises robustness of symmetry-breaking, while enhanced dimerization renders the network less responsive due to kinetic trapping of PAR-2 on internal membranes and reduced sensitivity of PAR-2 to membrane displacement by the polarity kinase, aPKC/PKC-3. Thus, our data reveal how a dynamically oligomeric RING domain results in a cell polarity network that is both robust and responsive and highlight how tuning of oligomerization kinetics can serve as a general strategy for optimizing dynamic and cooperative intracellular targeting.

## Introduction

Robust polarization of cells typically relies on feedback pathways to amplify and stabilize molecular asymmetries (Chau et al., 2012; Gierer and Meinhardt, 1972; Meinhardt and Gierer, 1974; Mogilner et al., 2012; Wedlich-Soldner et al., 2003). Critically, this feedback must be appropriately configured to balance key tradeoffs in the potential behaviors of a system. For example, increased feedback may render a system more sensitive to polarizing cues and enhance the stability of the resulting polarized state, but this may come at the cost of either responding to inappropriate cues such as random fluctuations or failing to adapt to signals that change in space and time (Jilkine and Edelstein-Keshet, 2011). While feedback is clearly implicated in the intracellular patterning mechanisms that underlie cell polarity, it is often difficult to obtain direct and quantitative measures of feedback in living systems, let alone be able to directly link feedback behavior to specific molecular activities such that feedback can be manipulated to test its effects on system behavior. This is due in part to inherent complexity and redundancy of polarity networks that make it difficult to isolate core feedback circuits and the technical challenge posed in performing the required dose-response measurements *in vivo* with sufficient precision and accuracy (Graziano et al., 2017). Thus, in many cases, we lack rigorous quantitative assessment of what are often purported to be core pattern-forming features of cell polarity networks.

The PAR (partitioning defective) polarity network is one such example. At the core of the PAR polarity network is a set of cross-inhibitory interactions that result in mutually exclusive localizations of distinct groups of peripherally associated PAR proteins on the plasma membrane (Goehring, 2014; Lang and Munro, 2017) (Figure 1A). In the *C. elegans* zygote, polarization is induced by cortical actomyosin flows that segregate one group of PAR proteins, the so-called aPARs that include PAR-3, PAR-6, and PKC-3 (aPKC), into an anterior membrane-associated domain (Goehring et al., 2011b; Munro et al., 2004). The anterior polarity kinase PKC-3 phosphorylates a second group of posterior polarity proteins (pPARs) that include PAR-1, PAR-2, and LGL-1 to restrict their localization: Prior to symmetry-breaking, pPAR proteins are initially depleted from the plasma membrane by aPARs and then load onto the posterior as aPARs are segregated by flows. (Betschinger et al., 2003; Hao et al., 2006; Hurov et al., 2004; Tabuse et al., 1998). The posterior polarity kinase PAR-1 in turn targets PAR-3, helping to restrict its localization to the anterior plasma membrane (Benton and St Johnston, 2003; Guo and Kemphues, 1995; Motegi et al., 2011). Polarity is further re-enforced by an additional reciprocal cross inhibitory circuit involving active CDC-42/PKC-3 in the anterior and the CDC-42 GAP, CHIN-1, in the posterior (Kumfer et al., 2010; Sailer et al., 2015).

**Figure 1.**
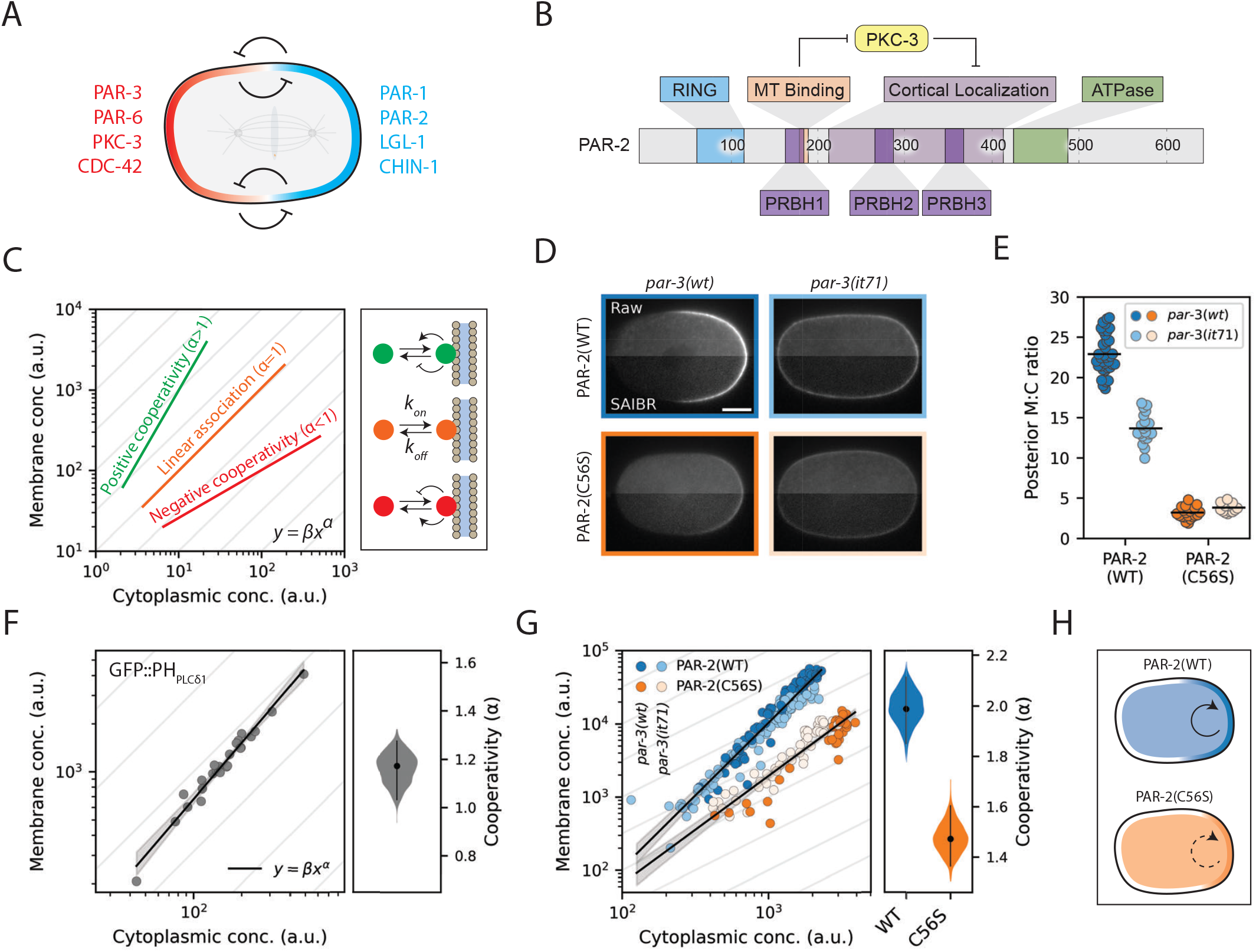
The PAR-2 RING domain drives cooperative membrane association. **(A)** PAR polarity relies on mutual antagonism between aPARs (red) and pPARs (blue) to maintain distinct anterior and posterior domains. **(B)** Schematic of PAR-2 functional domains. **(C)** Cooperative membrane association causes a deviation from linearity in the mapping between cytoplasmic and membrane concentrations. A molecule that binds and unbinds from the membrane with linear kinetics (i.e. rates independent of concentrations) will have a linear relationship between cytoplasmic and membrane concentrations, equivalent to a slope of 1 on a log-log plot (orange). Cooperative membrane association, in which the rates of membrane association and/or dissociation change as a function of membrane concentration, leads to a nonlinear mapping between cytoplasmic and membrane concentrations, equivalent to a slope *>*1 on a log-log plot in the case of positive cooperativity (green) or *<*1 in the case of negative cooperativity (red) (see Methods -Scoring cooperativity). **(D)** Raw and SAIBR-corrected images of mNG::PAR-2(WT) and mNG::PAR-2(C56S) in polarized (*par-3(WT)*) and uniform (*par-3(it71)*) conditions. Scale bar = 10 μm. **(E)** Quantification of posterior membrane to cytoplasmic ratio for the conditions in (D). **(F)** Quantification of membrane and cytoplasmic GFP::PH_PLC*8*1_ concentrations in cells with varying total amounts of GFP::PH_PLC*8*1_ (achieved by RNAi). Black line shows linear fit to log-transformed data and 95% confidence band. Right: probability distribution of the cooperativity score (slope of the linear fit), calculated by bootstrapping. A cooperativity score close to 1 reveals near-linear membrane association. Black vertical line shows 95% confidence interval. **(G)** Quantification of membrane and cytoplasmic PAR-2 concentrations in cells with varying total amounts of PAR-2. Data from both polarized cells (dark points) and uniform cells (light points) are pooled, for both wild type PAR-2 (blue) and RING-mutant (C56S) PAR-2 (orange). Membrane concentration measurements are limited to the posterior-most 20% of the cell. Black lines show linear fits to log-transformed data and 95% confidence bands. Right: probability distributions of the cooperativity scores for PAR-2(WT) and PAR-2(C56S). Black vertical lines show 95% confidence bands. **(H)** Schematic of PAR-2 cooperative membrane association. Posterior membrane concentrations of wild-type PAR-2 are amplified by cooperativity. Cooperativity is diminished in RING-mutant (C56S) cells, resulting in reduced membrane concentrations.

At the same time, it is increasingly thought that simple cross-inhibitory reactions are insufficient to fully account for the behavior of the PAR network, most notably because pattern formation by the PAR network, like many other patterning networks, is thought to depend on non-linear or bistable reaction dynamics that allow the system to support opposing membrane domains in distinct states (Arata et al., 2016; Dawes and Munro, 2011; Goehring et al., 2011b; Jilkine and Edelstein-Keshet, 2011; Lang and Munro, 2017; Meinhardt, 1982; Sailer et al., 2015). How such nonlinearity arises in this system remains unclear. While a number of mechanisms have been postulated, including a potential role for oligomerization (Arata et al., 2016; Dawes and Munro, 2011; Goehring et al., 2011b; Sailer et al., 2015), direct measurements of nonlinear feedback are generally lacking. Thus, the key links between molecular activities, feedback responses, and network behavior remain poorly explored.

Here we focus on a subsystem of the PAR network centered on the posterior PAR protein PAR-2 (Figure 1B). PAR-2 reversibly associates with the plasma membrane via a series of PRBH (**P**KC **R**esponsive **B**asic **H**ydrophobic) motifs that mediate electrostatic interaction with negatively charged lipids at the plasma membrane and which are thought to be targeted for phosphorylation by PKC-3 to induce membrane dissociation (Hao et al., 2006; Motegi et al., 2011). PAR-2 is not thought to directly antagonize anterior PAR proteins, but rather supports polarity through what is known as the eponymous PAR-2 pathway (Ramanujam et al., 2018; Zonies et al., 2010). In this proposed pathway, binding of PAR-2 to centrosomal microtubules allows it to locally avoid phosphorylation by PKC-3 in the posterior at the time of symmetry-breaking (Motegi et al., 2011). Once at the membrane, PAR-2 is thought to promote its own recruitment, and becomes stabilized against the action of PKC-3 via its RING (**R**eally **I**nteresting **N**ew **G**ene) domain (Arata et al., 2016; Hao et al., 2006; Motegi et al., 2011). PAR-2 in turn promotes recruitment of PAR-1 to the plasma membrane to support exclusion of PAR-3 from the posterior (Boyd et al., 1996; Motegi et al., 2011; Ramanujam et al., 2018). Normally this pathway re-enforces the aPAR asymmetry induced by flows. However, if actomyosin flows are disrupted and the initial segregation of aPARs fails, the PAR-2 pathway is sufficient to generate a posterior PAR-2 domain, which is stable despite initially overlapping with aPARs (Goehring et al., 2011b; Motegi et al., 2011; Zonies et al., 2010). Once formed, this domain drives clearance of aPARs from the posterior to establish a properly polarized zygote (Motegi et al., 2011).

The apparent ability of PAR-2 to self-organize into a membrane-associated domain despite a lack of obvious spatial input from anterior PAR proteins suggested to us that it may possess intrinsic self-amplifying feedback. We therefore set out to identify and define the nature of this feedback, and quantitatively link it back to the molecular properties of PAR-2.

## Results

### PAR-2 exhibits RING domain-dependent positive feedback

As a first step, we sought to determine whether PAR-2 exhibits cooperative membrane binding. A simple model of reversible membrane binding would be expected to yield a linear relationship between membrane and cytoplasmic concentrations with the membrane to cytoplasm (M:C) ratio given by *k*_on_/*k*_off_, where *k*_on_ and *k*_off_ define the respective membrane association and dissociation rate constants (Figure 1C). By contrast, in systems with positive and/or negative cooperativity, M:C ratios will be concentration-dependent. Typically, membrane-bound PAR-2 is present at higher concentrations when it is segregated within a posterior domain than when it is uniform (Figure 1D, (Cuenca et al., 2003; Hao et al., 2006)). While one might expect such an increase in posterior membrane concentration due to restriction of membrane-associated PAR-2 to a reduced area (i.e. PAR-2 is excluded from the anterior by PKC-3), if membrane binding is governed by mass action, M:C ratios should remain constant and thus be independent of whether PAR-2 is polarized. Thus, simply measuring membrane:cytoplasmic (M:C) ratios for PAR-2 and PAR-2 variants under polarized and uniform conditions should provide insight into whether membrane binding is concentration-dependent.

To accurately measure M:C ratios, we combined autofluorescence correction via SAIBR (Rodrigues et al., 2022) with a machine learning approach to assign local membrane and cytoplasmic fluorescence signals (Figure S1). Strikingly, M:C ratios for PAR-2 were increased nearly two-fold when PAR-2 was restricted to the posterior domain compared to when it was uniformly distributed (Figure 1D, 1E). This observation argues against a simple mass action model for membrane association of PAR-2.

We next sought to identify which features of PAR-2 were responsible for these polarization-dependent changes in membrane association. PAR-2 consists of an N-terminal RING domain, a region implicated in microtubule binding, a generally unstructured region enriched in basic-hydrophobic stretches that is required for membrane/cortex association, and a C-terminal ATPase domain which appears dispensable for function (Fig. 1B)(Hao et al., 2006; Levitan et al., 1994; Motegi et al., 2011). Mutations affecting the ATPase domain or microtubule-binding regions have shown minimal effects on membrane localization under normal conditions (Hao et al., 2006; Motegi et al., 2011). By contrast, while the N-terminal domain of PAR-2 has been reported to be insufficient for membrane association, variants of PAR-2 that either lack the RING domain or in which the RING is disrupted by mutation of a Zn-coordinating cysteine (C56S) exhibit reduced membrane association (Hao et al., 2006). Thus, while the RING domain appears to lack intrinsic membrane binding activity, it is required to potentiate membrane binding activity present elsewhere in the protein. We therefore introduced the C56S mutation into the *par-2* locus and measured the M:C ratio of PAR-2(C56S) in polarized and uniform conditions. In contrast to PAR-2(WT), PAR-2(C56S) exhibited M:C ratios that were similar between the segregated and uniform states (Figure 1D, 1E). Thus the RING domain of PAR-2 appears to be important for the apparent cooperativity in PAR-2 membrane binding.

To explicitly measure the degree of positive feedback in PAR-2 membrane association, we quantified the relationship between membrane and cytoplasmic concentrations in embryos subject to progressive reduction in total PAR-2 by RNAi. As a control, we examined embryos expressing a GFP fusion to the PIP_2_-binding domain of PLCo1 (GFP::PH_PLCo1_), which we could progressively deplete by *gfp(RNAi)*. Fitting of membrane to cytoplasmic concentrations with a simple cooperative binding model yielded an effective exponent, a, of less than 1.2, consistent with minimal cooperativity (Fig. 1F). By contrast, applying our method to embryos expressing endogenously-tagged mNG::PAR-2 that were subject to progressive depletion of PAR-2 by *par-2(RNAi)* yielded a ∼ 2, consistent with the existence of positive cooperativity (Fig. 1G). We obtained similar data regardless of whether we performed measurements at the posterior pole, where aPAR levels are low, and in a *par-3(it71)* mutant, in which PAR-2 intrinsic behavior is isolated from feedback from aPAR proteins (Figure 1G, Figure S2). Finally, consistent with a role for the RING domain in driving cooperativity, introduction of the RING-disrupting mutation C56S significantly reduced apparent cooperativity (Figure 1G).

We conclude that the RING domain drives effective cooperative membrane association of PAR-2 and that this cooperativity amplifies the ability of PAR-2 to be concentrated on the posterior membrane.

### RING domain dimerization is required for positive feedback

One mechanism to generate cooperativity would be for the PAR-2 RING domain to promote its own recruitment to the plasma membrane. To test whether the RING domain was sufficient to mediate such interactions, we expressed a soluble form of the RING domain and asked whether it could be recruited by endogenous PAR-2 to the posterior PAR domain. We found that an mNG::RING domain fusion was not recruited to the membrane, appearing identical to mNG alone (Figure 2A). This result was consistent with prior work showing that an N-terminal fragment containing the RING domain, but lacking predicted PRBH domains 2 and 3, failed to localize to the plasma membrane (Hao et al., 2006). However, when we tethered mNG::RING to membrane via fusion to PH_PLCo1_ (Audhya et al., 2005; Hurley and Meyer, 2001), it was efficiently recruited into the posterior PAR domain in a manner that depended on both an intact RING domain and the presence of endogenous PAR-2 (Figure 2B, C). Thus, the RING domain appears to be sufficient to mediate recruitment by endogenous PAR-2, provided that it is stabilized at the plasma membrane.

**Figure 2.**
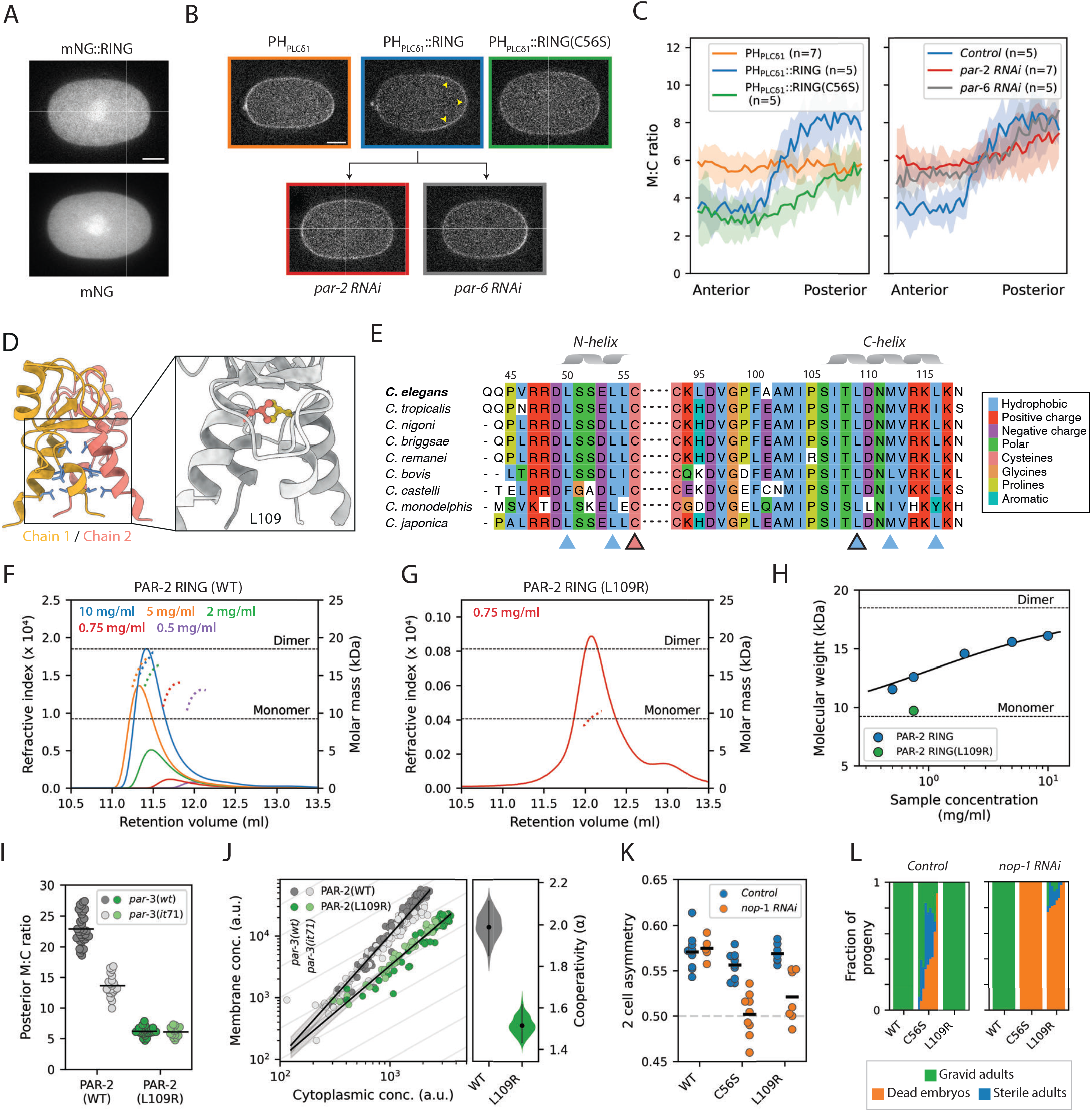
Cooperative membrane association arises from dimerization of the PAR-2 RING domain. **(A)** Isolated RING domain fragment displays no membrane association. SAIBR-corrected images of an mNG-tagged PAR-2 RING domain fragment, compared to mNG alone. Scale bar = 10 μm. **(B)-(C)** A membrane-tethered RING domain fragment displays posterior enrichment in a polarity-dependent manner. (B) SAIBR-corrected images of GFP::PH::RING compared to GFP::PH and GFP::PH::RING(C56S). Unlike GFP::PH and GFP::PH::RING(C56S), GFP::PH::RING displays considerable enrichment in the posterior (arrowheads), presumably through an interaction with endogenous PAR-2, which is lost upon disruption of underlying PAR polarity by either *par-2* or *par-6 RNAi*. Scale bar = 10 μm. (C) Anterior to posterior quantification of local membrane to cytoplasmic ratio for the lines and conditions in (B). Mean*±*SD. **(D)** AlphaFold structure prediction for the PAR-2 RING domain dimer (PAR-2 residues 40-120), with inward facing hydrophobics indicated (blue). Inset shows enlarged view of the 4-helix bundle with L109 indicated. **(E)** Clustal Omega alignments of the PAR-2 RING domain C and N helices within the *Caenorhabditis* genus. Arrowheads indicate inward-facing hydrophobic residues within the C and N helices (blue), including *C. elegans* L109 (black border). The zinc-coordinating residue C56 is also indicated (pink). **(F)** SEC-MALS traces for the PAR-2 RING domain at different sample concentrations. Solid lines indicate refractive index measurements, dotted lines indicate molar mass measurements. Color coded by sample concentration. **(G)** SEC-MALS trace for PAR-2 RING (L109R) at a sample concentration of 0.75 mg/ml. **(H)** The PAR-2 RING domain displays concentration-dependent dimerization. Average molecular weight measurements vs. sample concentration for the SEC-MALS assays in (F) and (G). Solid line indicates fit of wild type data to a dimer model (see Methods). **(I)** Quantification of PAR-2(L109R) posterior membrane to cytoplasmic ratio in polarized (*par-3(WT)*) and uniform (*par-3(it71)*) conditions. Wild type data from Figure 1 repeated in grey for reference. **(J)** Quantification of membrane and cytoplasmic concentrations of PAR-2(L109R) in cells with varying total amounts of PAR-2 (wild type data from Figure 1 repeated in grey for reference). Data from both polarized cells (dark points) and uniform cells (light points) are pooled. Right: probability distribution of the cooperativity score calculated by bootstrapping. Black vertical lines show 95% confidence internal. **(K)** Two-cell asymmetry (Area^AB^ / Area^AB+P1^) in wild type, *par-2(C56S)* and *par-2(L109R)* cells (control vs *nop-1 RNAi*). **(L)** Fraction of gravid adults, sterile adults and dead embryos in the progeny of wild type, *par-2(C56S)* and *par-2(L109R)* worms (control vs *nop-1 RNAi*). Each full bar is a composite of 8-10 smaller bars representing individual trials for each condition.

How then could the RING domain facilitate its own recruitment? The PAR-2 RING domain sequence harbors a C3HC4 pattern of zinc-coordinating residues that is characteristic of RING-family E3 ligases. Structural homology modeling of the PAR-2 RING domain suggested similarity to dimeric E3 ligases (https://swissmodel.expasy.org/) and an AlphaFold structure prediction for a PAR-2 RING dimer was similar to dimeric E3 RING domains, which are characterized by a four helix bundle consisting of an N and C helix from each of the two monomers (Figure 2D)(Jumper et al., 2021). The PAR-2 RING exhibits the expected “knobs-into-holes” pattern of conserved hydrophobic residues (L50, L54, L109, M112, L116) within the predicted four-helix bundle, mutation of which has been shown to disrupt dimerization of other RING domains (Figure S3)(Brzovic et al., 2001; Crick, 1953; Fiorentini et al., 2020) and which are broadly conserved within the Caenorhabditis genus (Figure 2E). As we were unable to demonstrate E3 ligase activity (data not shown) and given previous reports of PAR-2 oligomerization (Arata et al., 2016; Motegi et al., 2011), we wondered whether dimerization of the PAR-2 RING domain could underlie the cooperative membrane recruitment that we observe.

To test whether the PAR-2 RING domain was capable of oligomerization, we purified the PAR-2 RING domain from *E. coli* and subjected the purified RING domain to SEC-MALS to determine its oligomeric state. These data revealed concentration-dependent dimerization shifting from mostly monomeric to mostly dimeric over the concentration ranges tested (Figure 2F, H). To selectively disrupt dimerization, we mutated L109, the sidechain of which lies at the heart of the hydrophobic core of the putative dimer interface, making symmetric contact with L109 from the second protomer (Figure 2D). Consistent with predictions, the L109R RING domain was predominantly monomeric (Figure 2G, H).

Having established that L109R disrupts dimerization in vitro, we examined the effects of L109R in vivo. Quantification of membrane binding revealed that L109R reduced the M:C ratio nearly as much as C56S. L109R also showed similar M:C ratios between the polarized and uniform states and substantially reduced nonlinearity in the relationship between membrane and cytoplasmic concentrations, suggesting that disruption of the dimer interface weakens positive feedback (Figure 2I, J). We also tested the effects of an additional predicted interface mutation (L50R). L50R also reduced membrane binding, though to a lesser extent than L109R, and did not show any additive effects with L109R (Figure S4).

Phenotypically, L109R mutants did not exhibit any developmental defects under otherwise wild-type conditions, and thus did not fully phenocopy C56S, which showed significant levels of maternal effect embryonic lethality and sterility, consistent with improper germline specification (Figure 2L). This may be due to destabilization of the RING domain by the C56S, which could explain the reduced membrane affinity and somewhat lower overall protein amounts of C56S vs L109R (Table S2). Nonetheless, both alleles exhibited similar maternal effect embryonic lethality in a *nop-1(RNAi)* background in which symmetry-breaking was rendered dependent on the PAR-2 pathway due to a reduction in cortical flows. In our conditions, 100% of *nop-1(RNAi)* embryos exhibited normal development and gave rise to fertile adults, consistent with the semi-redundant contributions of cortical flow and the PAR-2 pathway to polarization (Rose et al., 1995; Tse et al., 2012; Zonies et al., 2010). By contrast, and consistent with RING mutants exhibiting defects in the PAR-2 pathway, combining *nop-1(RNAi)* with *par-2(L109R)* or *par-2(C56S)* resulted in a reduction in division asymmetry (Figure 2K) and >80% and 100% embryonic lethality, respectively (Figure 2L).

We therefore conclude that dimerization of the PAR-2 RING domain underlies concentration-dependent membrane binding, which is required for symmetry-breaking when cortical flows are compromised and embryos rely on the PAR-2 pathway. The more penetrant phenotype of C56S vs. L109R is not unexpected given the potential destabilizing effects of disrupting RING domain folding compared to selective targeting of the dimer interface. Consistent with this interpretation, attempts to purify the PAR-2(C56S) RING domain failed to yield usable quantities of intact protein (data not shown).

### A simple thermodynamic model of dimerization is sufficient to generate positive feedback

To understand how dimerization of PAR-2 generates positive feedback, we formulated a thermodynamic model based on dimerization of a reversibly bound membrane-associated molecule. We let molecules exist in one of four states, cytoplasmic monomer, cytoplasmic dimer, membrane monomer, and membrane dimer, the relative concentrations of which will depend on the strengths of dimerization and membrane association. Note that this model relies only on the assumptions that the molecule undergoes reversible dimerization, that dimers and monomers can reversibly associate with the membrane, and that these activities occur independently (Figure 3A, Supplemental Model Description).

**Figure 3.**
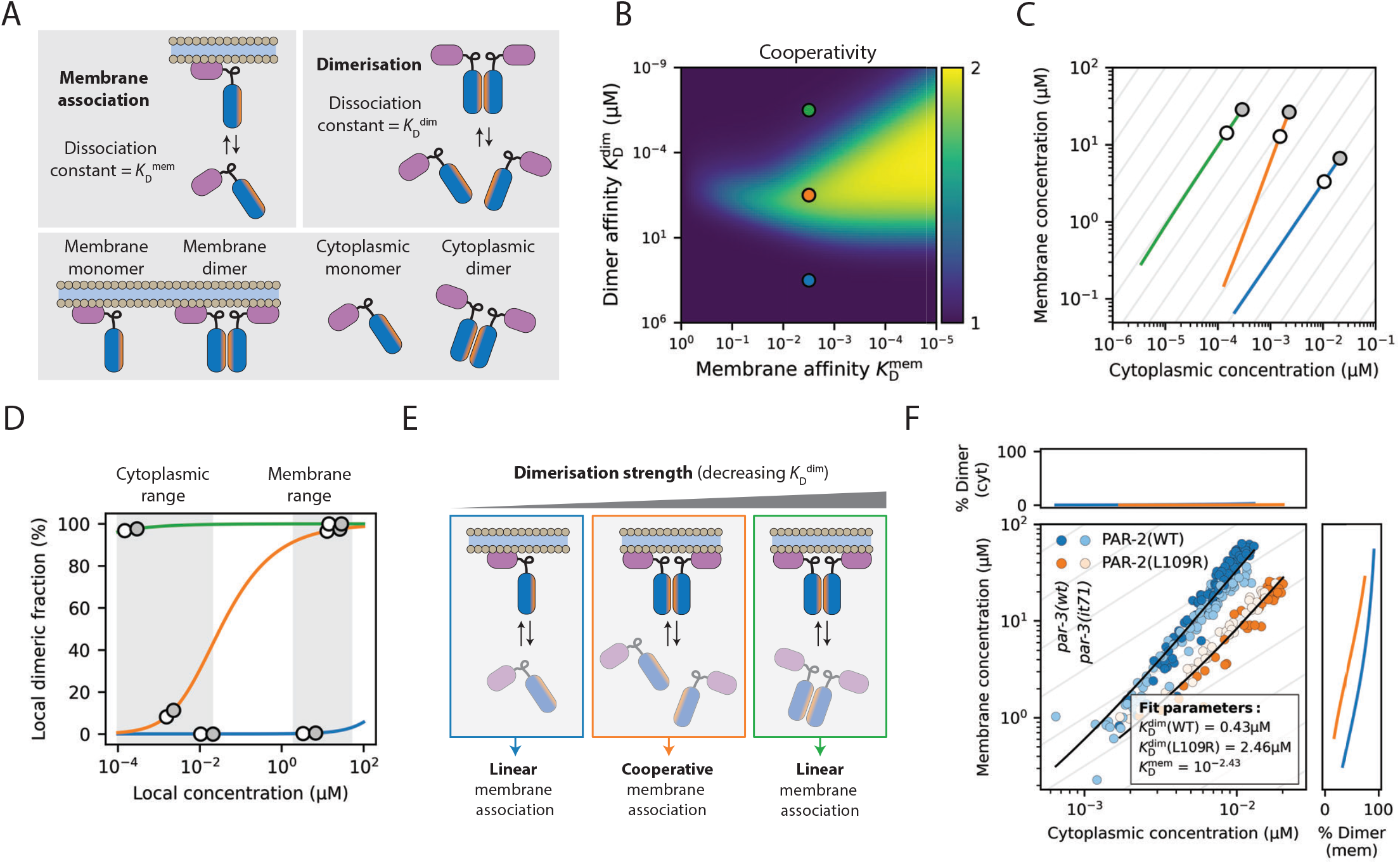
Cooperativity arises from selective stabilization of dimers at the plasma membrane. **(A)** Schematic for model of dimerization and membrane association. The model has two dissociation constants 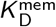 and 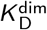 representing the strength ofmembrane association and dimerization respectively. **(B)** Membrane binding cooperativity as a function of dissociation constants 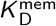 and 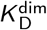, showing that cooperativity is optimized at intermediate dimerization strengths. Points in blue, orange and green correspond to parameter regimes shown in (C). **(C)** Mapping between cytoplasmic and membrane concentrations in systems with varying levels of total protein for the three parameter regimes indicated in (B). Grey and white points indicate systems with 100% and 50% total protein respectively. **(D)** Degree of dimerization as a function of local concentration for protein with high (green), intermediate (orange) and low (blue) dimerization strengths (corresponding to the 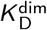 values shown in B). Local cytoplasmic and membrane concentrations corresponding to the grey and white points in (C) are shown for reference. **(E)** Cooperativity arises from selective stabilization of dimers at the membrane. Where dimerization strength is intermediate, protein exists largely as monomers in the weakly concentrated cytoplasm, but is induced to dimerize upon membrane association through an increase in local concentration, which further stabilizes membrane association. Systems in which dimer affnity is too low or too high are not induced to change dimeric state upon membrane binding, so membrane association is linear. **(F)** Fit of measured membrane vs cytoplasm relationships for PAR-2(WT) and PAR-2(L109R) to a model with shared 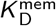 and different 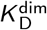 for WT vs L109R. Black lines show the best model fit. Panels top and right show degree of dimerization in the model as a function of local concentration across the relevant range of cytoplasmic and membrane concentrations.

Varying membrane binding (*K*_D_^mem^) and dimerization (*K*_D_^dim^) affinities revealed that cooperativity was maximal for high membrane binding affinity (low *K*_D_^mem^) and intermediate *K* _D_^dim^ (Figure 3B, 3C). This region of parameter space corresponded to a regime in which the dimer fraction was high at the membrane, but effectively absent in the cytoplasm (Figure 3D, 3E). Under such conditions, increases in local membrane concentration will stabilize membrane association via promoting dimerization.

Fits of this model to experimental RNAi rundown data for PAR-2(WT) showed good concordance. Specifically, we find that the estimated dimer affinity from the model fitting (*K*_D_^dim^ ∼ 425 nM, 95% CI [280, 654], Figure 3F, Table S4) reproduces the affinity that was independently measured in vitro by analytical ultracentrifugation (*K*_D_^dim^ (global fit) = 358 nM, Figure S5, Table S3). This was substantially higher than estimated cytoplasmic PAR-2 concentrations (10-50 nM)(Goehring et al., 2011b; Gross et al., 2019), consistent with PAR-2 being primarily monomeric in the cytoplasm and with our observation that only a membrane-tethered form of the isolated RING domain could be recruited by PAR-2 to the posterior PAR domain (Figure 2A-C). Simultaneous fitting of both PAR-2(WT) and PAR-2(L109R) with a common *K*_D_^mem^ indicate that L109R results in a ∼6-fold reduction in dimerization affinity (Figure 3F, Table S4). Constraining fits with measured values of *K*_D_^dim^ for PAR-2(WT) yielded similar results (Figure S6, Table S4).

Thus, a simple thermodynamic model of dimerization and membrane binding is sufficient to capture the cooperative membrane binding of PAR-2.

### Constitutive dimerization disrupts plasma membrane selectivity and PAR-2 function

A key prediction of our model is that both increasing or decreasing dimer affinity should compromise PAR-2 function. We have already shown that reduced dimer affinity compromised PAR-2 membrane recruitment and the robustness of polarization. To examine the effects of enhanced dimer affinity, we created a constitutive PAR-2 dimer by introducing a dimeric GCN4 leucine zipper at the end of the RING domain (40-120), before the first PRBH domains (Harbury et al., 1993; Illukkumbura et al., 2023). We found that PAR-2(GCN4) was enriched at the embryo posterior as PAR-2(WT), but showed residual membrane localization in the anterior membrane, suggesting it was less sensitive to removal by aPKC (Figure 4A, 4B). Unexpectedly, it also exhibited prominent accumulation on internal structures and a corresponding reduction in plasma membrane concentrations, suggesting that the normal preferential localization of PAR-2 to the plasma membrane is disrupted by constitutive dimerization (Figure 4Aii, iv). The enrichment of PAR-2(GCN4) near the centrosomes resembles known localizations of RAB-5/RAB-7/RAB-11, suggesting association with endosomal membranes (Hyenne et al., 2012; Zhang et al., 2008). This effect was not due to aberrant membrane targeting by the GCN4 sequence as an mNG::GCN4(dimer) fusion was diffusely localized in the cytoplasm (Figure 4C). Finally, to validate that this effect was not specific to GCN4, we introduced an alternative dimerization motif (6HNL, (Chang and Dickinson, 2022)), which yielded similar increases in residual anterior membrane localization and accumulation on internal membranes (Figure S7).

**Figure 4.**
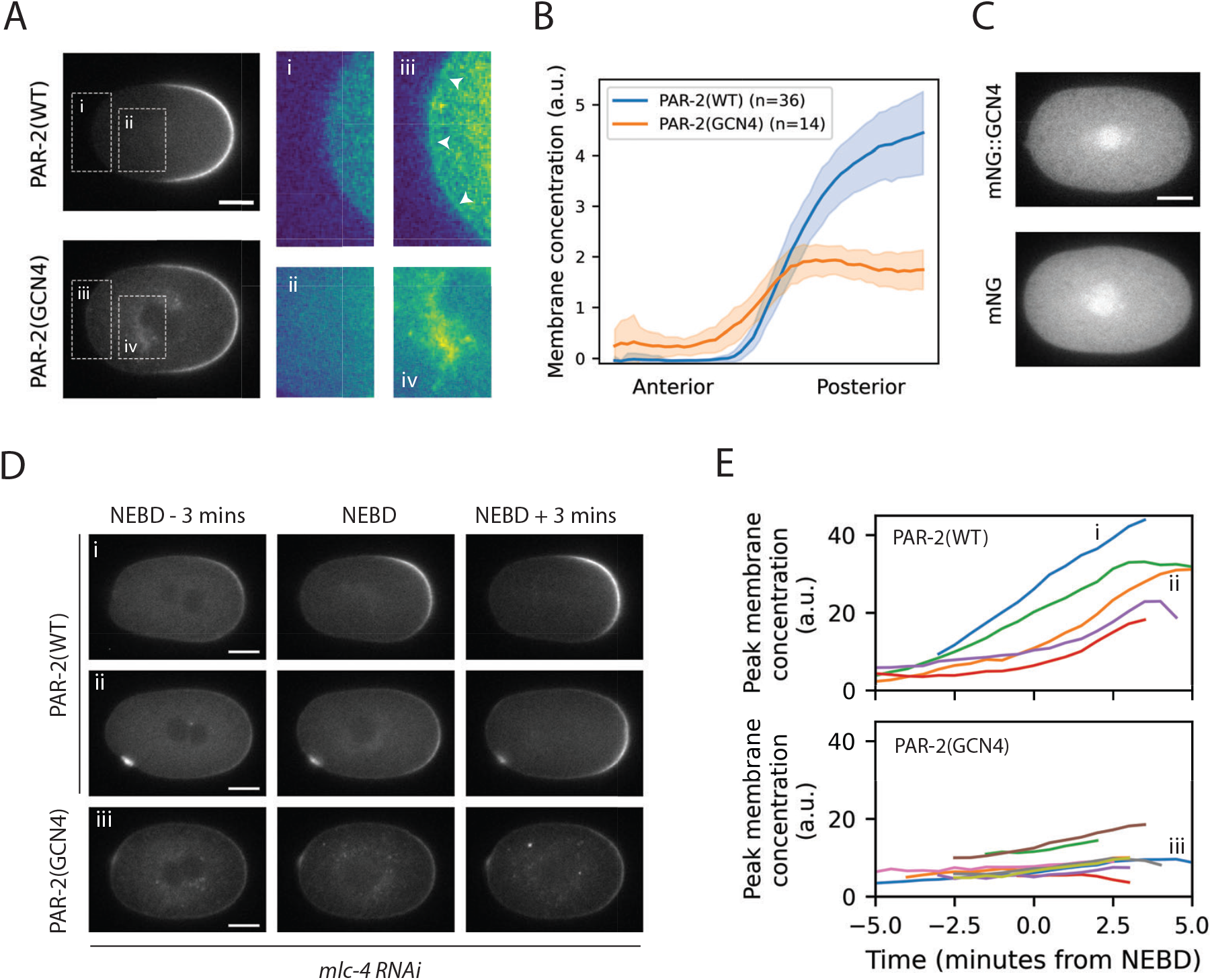
Enhanced dimerization leads to loss of membrane specificity and reduced sensitivity to PKC-3. **(A)** SAIBR-corrected images of PAR-2 and PAR-2(GCN4). Note the localization of PAR-2(GCN4) to the anterior membrane (iii, arrowheads), as well as prominent localization to internal structures (iv). Scale bar = 10 μm. **(B)** Anterior to posterior membrane concentration profiles of PAR-2(WT) and PAR-2(GCN4). Mean*±*SD. **(C)** GCN4 alone does not localize to the plasma membrane or internal structures. SAIBR-corrected image of mNG::GCN4 compared to mNG alone. Scale bar = 10 μm. **(D)-(E)** PAR-2(GCN4) displays defective symmetry breaking in *mlc-4 RNAi* conditions. (D) SAIBR-corrected images of PAR-2(WT) and PAR-2(GCN4) at three minutes pre-NEBD (nuclear envelope breakdown), NEBD and three minutes post-NEBD in *mlc-4 RNAi* conditions. Wild-type PAR-2 shows variation in the timing and degree of symmetry-breaking (i vs ii). Scale bars = 10 μm. (E) Quantification of peak membrane concentration over time for PAR-2(WT) and PAR-2(GCN4) in *mlc-4 RNAi* conditions. i, ii and iii correspond to the embryos in (D).

While *par-2(GCN4)* animals did not show significant lethality under normal conditions, when we blocked cortical flows via depletion of a myosin regulatory light chain using *mlc-4(RNAi)*, the efficiency of polarization was reduced, consistent with defects in the PAR-2 pathway (Figure 4D, 4E). While embryos were often capable of generating some asymmetry, PAR-2 domains were substantially less pronounced and were accompanied by significant levels of residual membrane-associated PAR-2 in the anterior. Thus, somewhat counter-intuitively, increasing dimerization strength reduces the ability of PAR-2 to be targeted to the posterior plasma membrane during polarization. Because both increasing and decreasing dimer affinity disrupts polarization under conditions in which the PAR-2 pathway is required, we conclude that PAR-2-dependent polarization relies on optimization of RING dimer affinity.

### Enhanced membrane affinity of ectopic PAR-2 dimers leads to kinetic trapping on inappropriate membranes

How can we explain the loss of plasma membrane specificity of PAR-2(GCN4)? Plasma membrane selectivity for proteins like PAR-2 that bind non-specifically to negatively charged lipids is generally thought to rely on the differential (i.e. more negative) charge profile of the plasma membrane relative to internal membranes (Yeung et al., 2008). We therefore hypothesized that constitutive dimerisation may provide a sufficient avidity enhancement to allow stable binding of PAR-2(GCN4) to internal membranes, despite a weaker charge profile on these membranes.

We therefore introduced a second internal membrane compartment to the model with a reduced binding affinity (*K*_D_ ^int^), reflecting the normal preference of PAR-2 for the plasma membrane. We found that increasing dimerization generally favors partitioning to membrane compartments, leading to increased concentrations on internal membranes, consistent with dimer-dependent stabilization (Figure 5A). However, we did not observe an enhancement of partitioning to internal membranes at the expense of plasma membrane targeting as we observed for PAR-2(GCN4) in vivo. Instead, the relative preference for the plasma membrane increased with dimerization affinity. Thus, from an equilibrium perspective, an increase in dimerization affinity cannot explain the observed decrease in plasma membrane selectivity.

**Figure 5.**
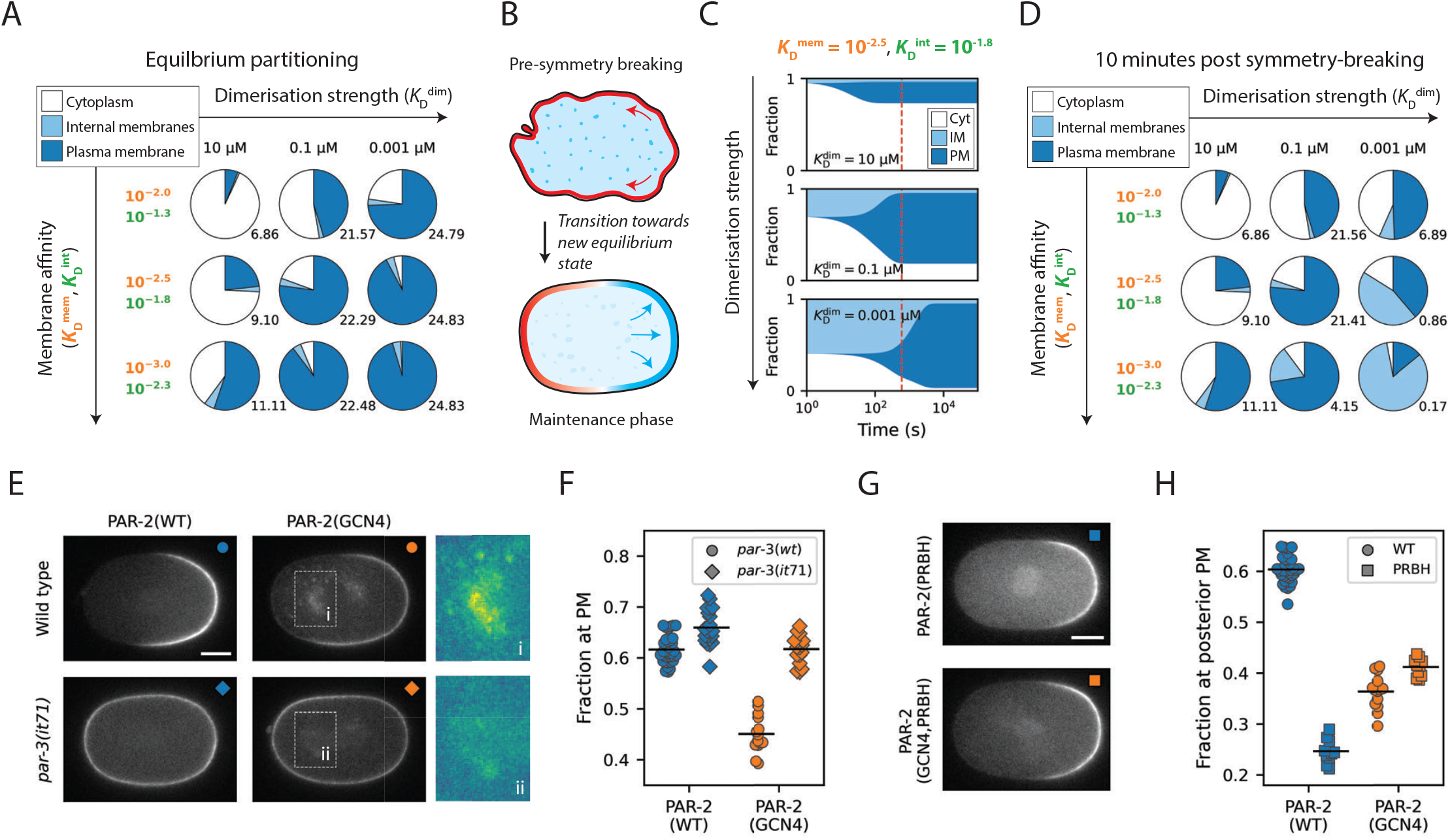
Intermediate dimerization affinity optimizes exchange kinetics to ensure membrane specificity. **(A)** Equilibrium solutions for a three-compartment model (cytoplasm, internal membranes (IM), and plasma membrane (PM)) as a function of dimerization strength 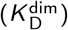 and membrane affnity (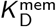 and 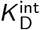). Pie charts show the fraction of total protein in each compartment at equilibrium. Numbers to the bottom right of each chart show the ratio of protein in the PM and IM compartments. 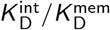 is fixed to a constant ratio of 5. **(B)** Schematic for symmetry breaking described as a transition between two equilibrium states. The system begins in a pre-symmetry breaking state in which only the cytoplasm and internal membrane compartments are available to PAR-2, with PAR-2 being held of the PM by the uniform activity of aPARs (red). At symmetry breaking, transport of aPARs to the anterior relieves inhibition of PAR-2 at the posterior plasma membrane, and the system transitions towards a new state reflecting this change in membrane availability. **(C)** Fraction of protein in each of the three compartments over time as the system transitions between a two compartment model (cytoplasm and IM only) and a three-compartment model (PM added), as a function of dimerization strength. When dimerization is weak (top), the system is fast to reach a new equilibrium, but with weak PM association. When dimerization is strong (bottom), the final equilibrium state has strong PM association, but the system takes exponentially longer to reach this state. With intermediate dimerization (middle), an intermediate behavior is observed. Dashed red line marks 600 seconds, roughly corresponding to the timing of NEBD after symmetry breaking. **(D)** PM specificity 10 minutes post symmetry breaking is maximized in intermediate dimerization regimes. As in (A), but showing a snapshot 10-minutes into the transition between the pre-symmetry breaking and maintenance phase states. Middle row corresponds to the simulations in (C). Note that in high dimerization regimes, PM association and specificity is maximized when membrane affnity is reduced as a result of increased transition kinetics. **(E)** SAIBR-corrected images of PAR-2 and PAR-2(GCN4) in *par-3(WT)* and *par-3(it71)* backgrounds. Note the reduced IM association of PAR-2(GCN4) in *par-3(it71)* conditions (ii vs i). Scale bar = 10 μm. **(F)** Fraction of total PAR-2 at the PM in each of the four conditions in (E). **(G)** SAIBR-corrected images of PAR-2(PRBH) and PAR-2(GCN4,PRBH). Note the lack of IM association in PAR-2(GCN4,PRBH) (compare to (E), top row). Scale bar = 10 μm. **(H)** Fraction of total protein at the posterior PM for PAR-2(WT), PAR-2(GCN4), PAR-2(PRBH) and PAR-2(GCN4,PRBH). Whereas mutating PRBH residues in PAR-2(WT) strongly decreases posterior plasma membrane association, adding these mutations to PAR-2(GCN4) leads to a moderate *increase* in posterior plasma membrane association.

One aspect of our system we have so far largely ignored is the relevant timescale of polarization, which is neglected in analysis of equilibrium conditions. During polarization, PAR-2 must shift from being nearly fully excluded from the plasma membrane by the activity of PKC-3 at the end of meiosis II to being enriched on the posterior membrane as PKC-3 is segregated into the anterior at the start of mitosis -a span of ∼ 10 minutes (Figure 5B)(Cuenca et al., 2003; Reich et al., 2019). These dynamics place temporal constraints on membrane binding -if membrane affinity is too high, redistribution of PAR-2 between membrane compartments may simply be too slow, leaving PAR-2 kinetically trapped on internal membranes. Consistent with this picture, we found that PAR-2(GCN4) exhibits both reduced mobility in the cell interior and slower redistribution from the cell interior onto the posterior plasma membrane during polarization (Figure S8).

To explore how dimerization affinity influenced polarization timescales in our model, we used transition state theory to assess the time evolution of our dual membrane system as it shifted from an unpolarized equilibrium state in which molecules only have access to the low affinity internal membrane compartment, and a polarized equilibrium state in which molecules gain access to the plasma membrane, reflecting the clearance of PKC-3 from the posterior membrane of the zygote during polarization. For all values of *K*_D_ ^dim^, molecules were initially excluded from the plasma membrane and then relocalized to the plasma membrane over time at the expense of both the cytoplasmic and internal membrane compartments. Importantly, all eventually reached a polarized state in which concentrations at the plasma membrane exceeded that on internal membranes, reflecting the differential affinities for the two membrane compartments (Figure 5C). However, as we suspected, given the stabilizing effect of dimerization on membrane binding, increasing dimerization strength dramatically slowed the timescale of this redistribution from internal pools to the plasma membrane in the model (Figure 5C, S9). Consequently, if assessed at intermediate timepoints (e.g. ∼10 minutes), increasing dimerization affinity appears to enhance the internal membrane pool at the expense of the plasma membrane (Figure 5D). Note, this apparent loss of selectivity arises purely from slower kinetics caused by dimerization-dependent reduction of membrane dissociation such that at similar time points, the strong dimer system is much further from the equilibrium, plasma-membrane dominated state.

This model therefore predicts that kinetic trapping of PAR-2 on internal membranes will be reduced if we either extend the time available for PAR-2 to equilibrate between the internal and plasma membrane compartments or compensate for the increase in avidity provided by enhanced dimerization by reducing the affinity of monomers for membranes. To increase the time available for equilibration, we examined the behavior of PAR-2(GCN4) in *par-3(it71)* embryos, which lack PKC-3 activity at the membrane and thus PAR-2 is not cleared from the membrane at the end of meiosis II (Reich et al., 2019; Tabuse et al., 1998). Consistent with predictions, PAR-2(GCN4) exhibited reduced levels of localization to internal membranes in *par-3(it71)* embryos compared to *par-3(wt)* embryos (Figure 5E, 5F).

To reduce membrane affinity of monomers, we targeted one of three putative PRBH motifs that are thought to mediate PAR-2 membrane association (Bailey and Prehoda, 2015; Brzeska et al., 2010; Ramanujam et al., 2018)(Figure 1B). Replacement of 7 serines to glutamic acid is sufficient to prevent PAR-2 enrichment at the plasma membrane (Hao et al., 2006). We therefore introduced two S>E mutations into the PRBH3 region (S334E, S338E) to achieve a modest reduction in membrane affinity. When introduced into the wild-type context, these mutations resulted in reduced accumulation within the posterior PAR domain, consistent with reduced membrane affinity (Figure 5G, 5H, PRBH). Strikingly, these same mutations had the opposite effect in the context of the constitutive dimer. PAR-2(PRBH,GCN4) exhibited enhanced accumulation within the posterior PAR domain (Figure 5H) and substantially reduced accumulation on internal membranes (Figure 5G) compared to PAR-2(GCN4). We also found that the introduction of PRBH mutations into PAR-2(GCN4) restored cytoplasmic turnover rates to near wild-type levels as well as sensitivity to PKC-3, with PAR-2(GCN4, PRBH) showing none of the residual anterior localization seen for PAR-2(GCN4) (Figure 5G, S8). Thus, as predicted by our model, one can at least partially rescue the effects of constitutive dimerization (decreased *K*_D_^dim^) by reducing the intrinsic membrane binding affinity of the constituent monomers (increased *K*_D_^mem^) (Figure 5D).

We therefore conclude that constitutive dimerization kinetically traps PAR-2 on internal membranes through enhanced membrane binding, severely increasing the timescale required for plasma membrane accumulation during polarization.

The fact that the robustness of polarization by the PAR-2 pathway is compromised by both increases and decreases in dimer affinity strongly suggests that dimerization affinity is optimized. Such optimisation ensures that membrane binding of PAR-2 is both sufficiently cooperative to drive robust polarity, but also sufficiently dynamic that PAR-2 remains highly responsive to spatiotemporal changes to the system, such as those involved in symmetry-breaking.

## Discussion

Here we have identified cooperative membrane binding of PAR-2 as a key feature of the PAR polarity network in *C. elegans* and directly linked this behavior to optimized dimerization of its RING domain.

Although the vast majority of RING domain-containing proteins identified in humans are believed to act as E3 ubiquitin ligases (Deshaies and Joazeiro, 2009), a key role of RING domains is to facilitate protein-protein interactions (Borden, 2000). In the case of E3 ligases, RING domains typically recruit E2 ubiquitin conjugating enzymes to facilitate substrate modification. Importantly for our work, E3 ligases often act as multimers in which RING-RING interactions play critical roles in mediating interactions between E3 monomers or in E2 recruitment (Fiorentini et al., 2020). While we cannot rule out that PAR-2 possesses E3 ubiquitin ligase activity, our data suggest that it is this dimerization function of the RING domain that is critical in defining the cooperative nature of PAR-2 membrane association.

Specifically, we show that membrane binding cooperativity emerges from the optimization of dimer affinity such that the *K*_D_ is intermediate between the effective cytoplasmic and membrane concentrations. Consistent with this model, both increasing or decreasing dimerization affinity impacted the ability of PAR-2 to polarize. RING domains of E3 ligases can exhibit a broad range of dimer affinities (Fiorentini et al., 2020) and, analogously to what we have shown here, differences in RING dimer affinity in E3 ligases have been proposed to underlie distinct modes of substrate binding and activity regulation (Koliopoulos et al., 2016). We therefore suggest that the RING domain provides a highly tunable platform for dimer optimization, a feature which appears in this case to have been co-opted to facilitate robust symmetry-breaking in the PAR polarity network.

Cooperativity does not arise from direct recruitment of cytoplasmic monomers by membrane associated species, which is negligible in this system due to the low concentration of cytoplasmic molecules, a conclusion supported by the failure of isolated RING domains to be recruited by PAR-2 in the posterior domain. Rather, effective positive feedback arises because local increases in membrane concentration will favor dimerization of membrane-associated monomers, which will in turn render them more stably associated with the membrane (Agudo-Canalejo et al., 2020). It is therefore specifically membrane-dependent dimerization that accounts for the observed positive feedback.

Reinforcing the need to optimize dimer affinity, increasing dimer affinity led to loss of plasma membrane selectivity. We initially considered that the impact of dimerization on nonlinear dynamics might lead to a reduction in the relative preference of PAR-2 for the plasma vs. internal membranes. However, if anything, increasing dimerization favored plasma membrane binding in our equilibrium model. Instead, we found that loss of selectivity was due to a kinetic mismatch between the timescales of polarization and membrane turnover of the constitutive dimer on membranes. Due to enhanced stabilization of membrane association by constitutive dimers, redistribution of PAR-2(GCN4) dimers from internal membrane pools to the plasma membrane at the onset of polarization is simply too slow. Even in the absence of large scale reorganization, such dynamic redistribution is likely to be required to counter internalization of membrane-associated molecules by endocytosis and may explain why we observe some level of internal membrane association even when PAR-2(GCN4) is rendered resistant to membrane displacement by aPARs (e.g. in *par-3* embryos).

It has been speculated that membrane-stabilized oligomeric assemblies can constitute an effective memory in polarizing systems (Illukkumbura et al., 2023; Lang and Munro, 2022). By effectively slowing the timescale of membrane turnover, oligomerization can amplify and lock in the effects of transient polarizing cues. However, our data suggest that this “memory” comes at the cost of reduced responsiveness of the system as stable dimers are slow to adapt to changes in cell state. Our work therefore highlights how optimization of oligomerization kinetics, in this case of a dimeric RING domain, allows systems to balance this trade-off between memory and responsiveness in a dynamic system, which in the case of the PAR network facilitates robust and timely polarity establishment. Notably, both reduction and enhancement of dimer affinity impair the response of PAR-2 to symmetry-breaking cues and lead to defects in the PAR-2-dependent polarization pathway. Given the widespread occurrence of dynamic oligomerization and oligomerization-dependent localization within molecular networks, including the PAR and other polarity networks (Benton and Johnston, 2003; Dodgson et al., 2013; Harris, 2017; Lang and Munro, 2022; Meca et al., 2019; Mizuno et al., 2003; Sailer et al., 2015; Strutt et al., 2011), this paradigm of optimized and reversible oligomerization kinetics is likely to be a broadly applicable strategy for rapid and cooperative intracellular targeting.

## Contributions

Conceptualization: T.B., D.Z., N.W.G.; Methodology: T.B., D.B., N.H., R.R., D.Z.; Formal analysis: T.B., D.B., R.R., K.N., D.Z.; Investigation: T.B., D.B., N.H., R.R., D.Z., N.W.G.; Resources: T.B., D.B., N.H.; Writing – original draft preparation: T.B., R.R., N.W.G.; Writing – review and editing: All the authors; Supervision: N.Q.M., D.Z., N.W.G.; Project administration: N.W.G.; Funding acquisition: N.Q.M., D.Z, N.W.G.

## Acknowledgements

We would like to thank Ian Taylor, Laura Masino, the Schreiber and Rittinger Labs, and the Structural Biology Scientific Technology Platform at the Crick for help in purification and analysis of PAR-2 and the Goehring Lab and Katrin Rittinger for comments on the manuscript. Some strains were provided by the Caenorhabditis Genome Center (CGC), which is funded by NIH Office of Research Infrastructure Programs (P40 OD010440). This work was supported by the Francis Crick Institute, which receives its core funding from Cancer Research UK (CC2119, CC2068), the UK Medical Research Council (CC2119, CC2068), and the Wellcome Trust (CC2119, CC2068). D.Z. and R.R. gratefully acknowledge funding from the Max Planck Society and the European Union (ERC, EmulSim, 101044662). For the purpose of Open Access, the author has applied a CC BY public copyright license to any Author Accepted Manuscript version arising from this submission.

## Data Availability

Source code and documentation will be made available at https://github.com/goehringlab. Original datasets will be provided upon request from the corresponding author.

## Supplemental Materials

- Supplemental Materials and Methods
- Supplemental Figures S1 -S9
- Supplemental Tables S1-S4

