## Supplemental Figures and Tables for "Optimized dimerization of the PAR-2 RING domain drives cooperative and selective membrane recruitment for robust feedback-driven cell polarization"

*Bland et al. (2023)*

### Supplemental Figures

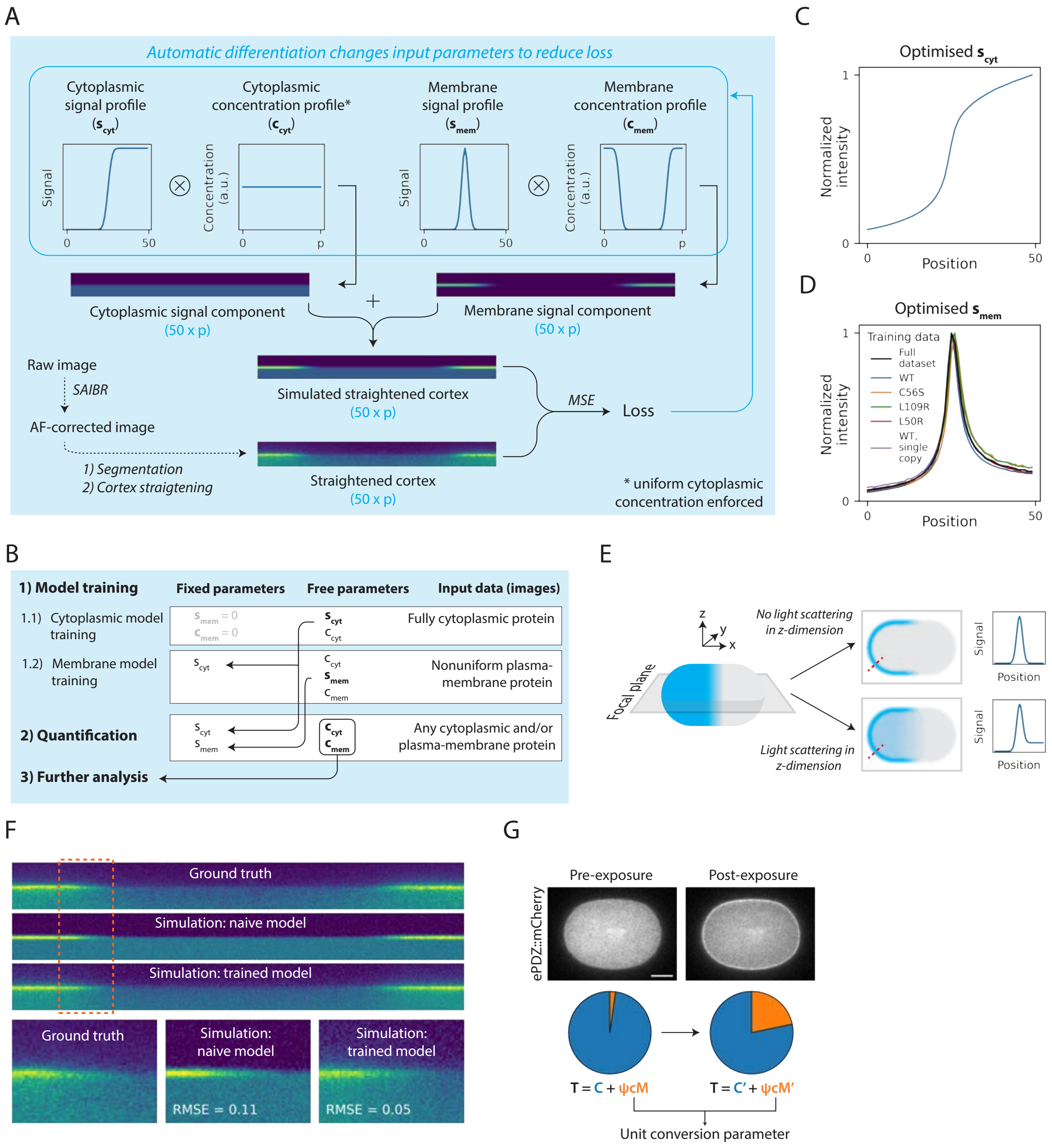

**Figure S1. A machine-learning method for extraction of normalized membrane and cytoplasmic protein concentrations from midplane confocal images. (A)** Schematic of differentiable model for image quantification. See Methods for details. **(B)** Outline of model training and quantification protocol. See Methods for details. **(C)** Cytoplasmic signal profile determined by cytoplasmic model training on images of cytoplasmic mNG. **(D)** Membrane signal profiles determined by membrane model training on images of wild type PAR-2, mutant alleles and single-mNG heterozygotes. Black line shows a model trained on the full dataset. **(E)** Schematic of the expected effects of 3D light scattering on observed midplane signal distributions from membrane protein. **(F)** Example of ground truth (SAIBR-corrected) and simulated images for an mNG::PAR-2(L109R) embryo. Naive model refers to a model in which membrane and cytoplasmic signal profiles are fixed to a Gaussian and error function. Trained model refers to a model in which cytoplasmic and membrane profiles have been trained according to the process outlined in (B). Gaussian noise has been added to simulated images to allow for closer visual comparison to the ground truth image. RMSE: root mean square error. **(G)** Optogenetics system used to calibrate cytoplasmic and membrane concentration units. Exposure to blue light promotes an interaction between ePDZ::mCherry and membrane-tethered PH::eGFP::LOV, causing recruitment of ePDZ::mCherry to the membrane. Pie charts show the amount of total ePDZ::mCherry in the cytoplasm and membrane before and after exposure to blue light, which sums to a constant value T. By solving the two equations shown, a unit conversion factor (c) can be calculated. Scale bar = $10\,\mu$m.

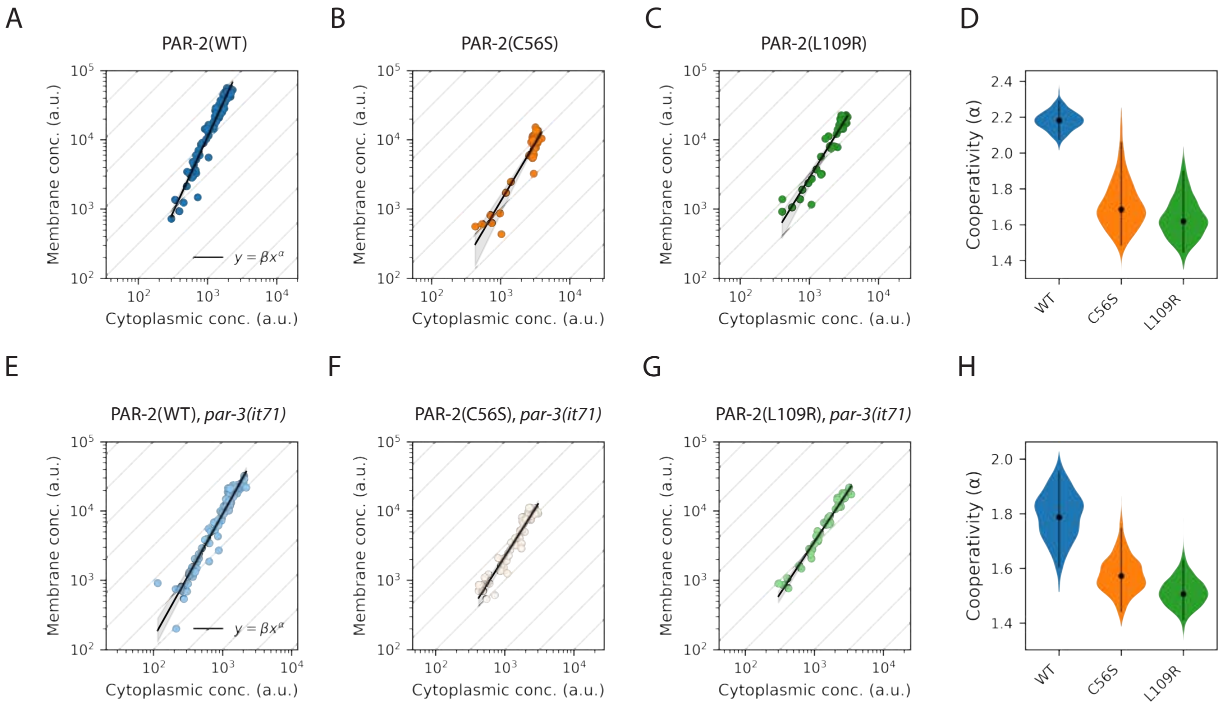

**Figure S2. PAR-2 cooperativity measurements in datasets stratified by polarity state. (A)-(C)** Membrane vs cytoplasmic concentrations in polarized (*par-3(WT)*) cells for wild type and RING mutant PAR-2 lines. Black lines show fits to a linear regression model and 95% confidence bands. **(D)** Probability distribution of cooperativity scores determined from polarized (*par-3(WT)*) data. Black vertical lines show 95% confidence intervals **(E)-(G)** Membrane vs cytoplasmic concentrations in uniform (*par-3(it71)*) cells. **(H)** Probability distribution of cooperativity scores determined from uniform (*par-3(it71)*) data.

A

| Protein | PDB code | Oligomeric state |
|---|---|---|
| TRIM69 | 6YXE | Dimer |
| RAD18 | 2Y43 | Dimer |
| RNF8 | 4AYC | Tetramer |
| TRIM21 | 6FGA | Dimer |
| TRIM25 | 5EYA | Dimer |
| TRIM5 | 4TKP | Monomer |

B

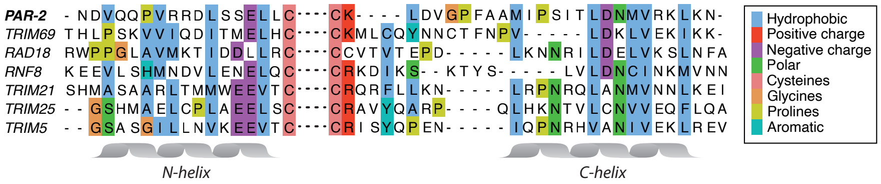

**Figure S3. The PAR-2 RING domain displays the characteristic pattern of hydrophobic residues expected for dimeric RING domains. (A)** RING domains in the Protein Data Bank (PDB) with closest homology to the PAR-2 RING domain (identified using the SWISS-MODEL homology modelling server). **(B)** Clustal Omega alignments of the PAR-2 RING domain N and C helices against the RING domains in (A). Note the characteristic pattern of hydrophobic residues (blue).

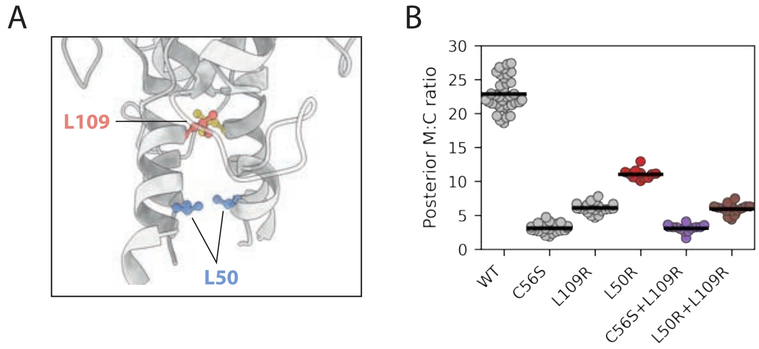

**Figure S4. Additional PAR-2 RING mutant alleles. (A)** PAR-2 RING domain dimer structure prediction (AlphaFold) with residues L50 and L109 highlighted. **(B)** Quantification of posterior membrane to cytoplasmic ratio for wild type and RING mutant PAR-2. Wild type, C56S and L109R data (grey) repeated from Figures 1 and 2 for comparison.

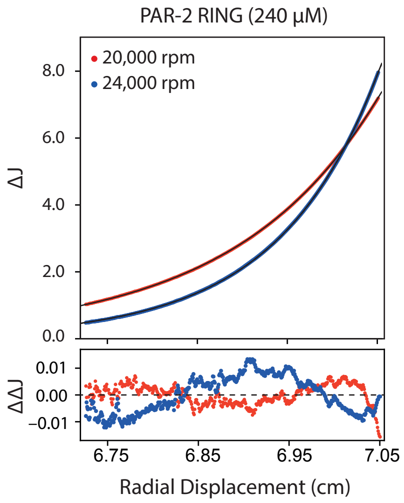

**Figure S5. PAR-2 RING self-associates in solution.** Multi-speed sedimentation equilibrium profiles determined from interference data collected on PAR-2 RING at 240 µM. Data was recorded at the speeds indicated. The solid black lines represent the global best fit to the data (red and blue points) using a monomer-dimer model ($K_D^{dim} = 0.36$ µM, reduced $\chi^2 = 1.25$). The lower panel shows the residuals to the fit.

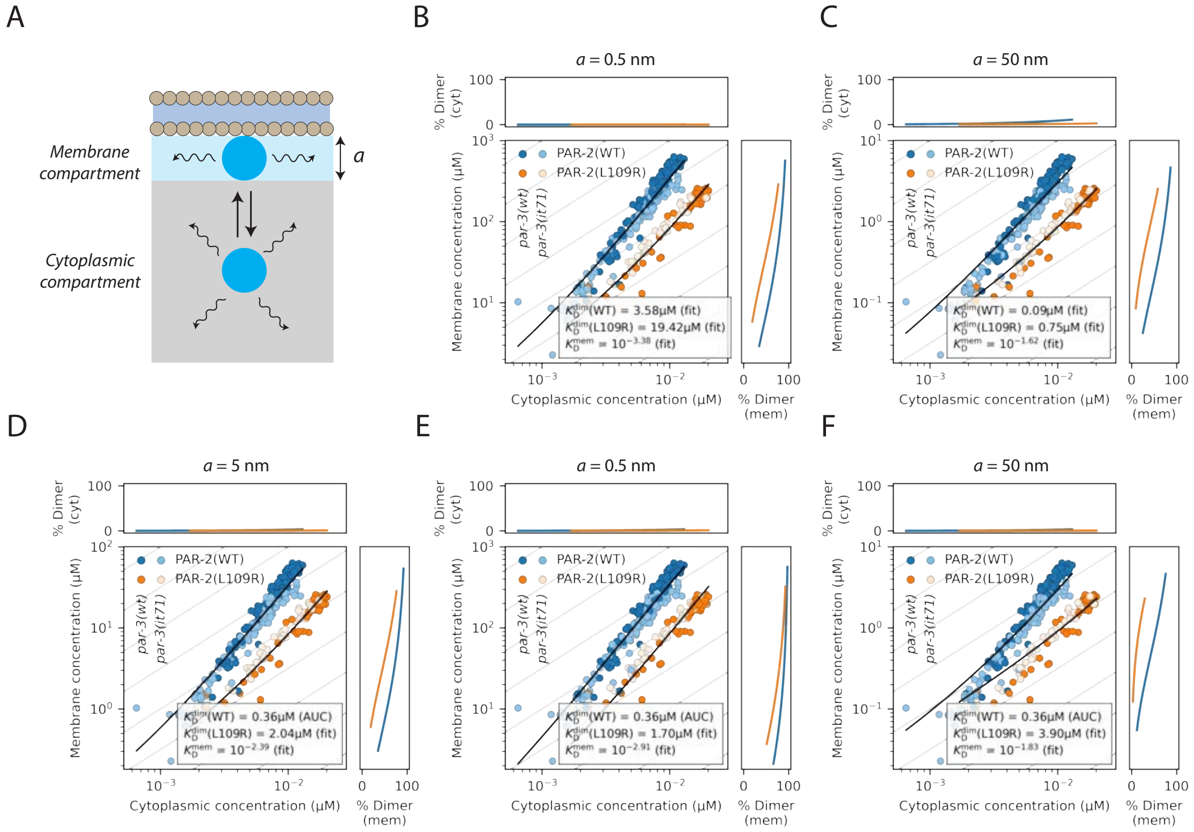

**Figure S6. Modelling the plasma membrane as a thin volume compartment. (A)** Schematic of the equilibrium model for membrane association. The system can be described as two volume compartments representing cytoplasmic and membrane-bound states. We assume that diffusion on the membrane compartment is confined to the two-dimensional plane of the plasma membrane, so the membrane compartment can be described as a volume with thickness $a$, where $a$ is equal to the diameter of the molecule. **(B)** - **(C)** Model fits as in Figure 3F, but using a value for $a$ 10x smaller (B) or 10x larger (C). **(D)** - **(F)** Model fits for three values of $a$ with $K_D^{dim}$ (WT) fixed to the value experimentally determined by AUC.

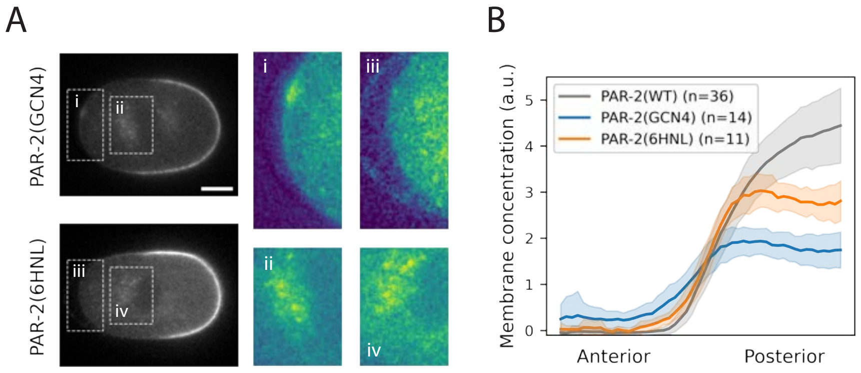

**Figure S7. Alternative dimerization domain (6HNL) yields similar phenotype to GCN4. (A)** SAIBR-corrected images of PAR-2(GCN4) and PAR-2(6HNL). Scale bar = $10\,\mu m$. **(B)** Anterior to posterior membrane concentration profiles of PAR-2(GCN4) and PAR-2(6HNL), with PAR-2(WT) shown for reference. Mean $\pm$ SD.

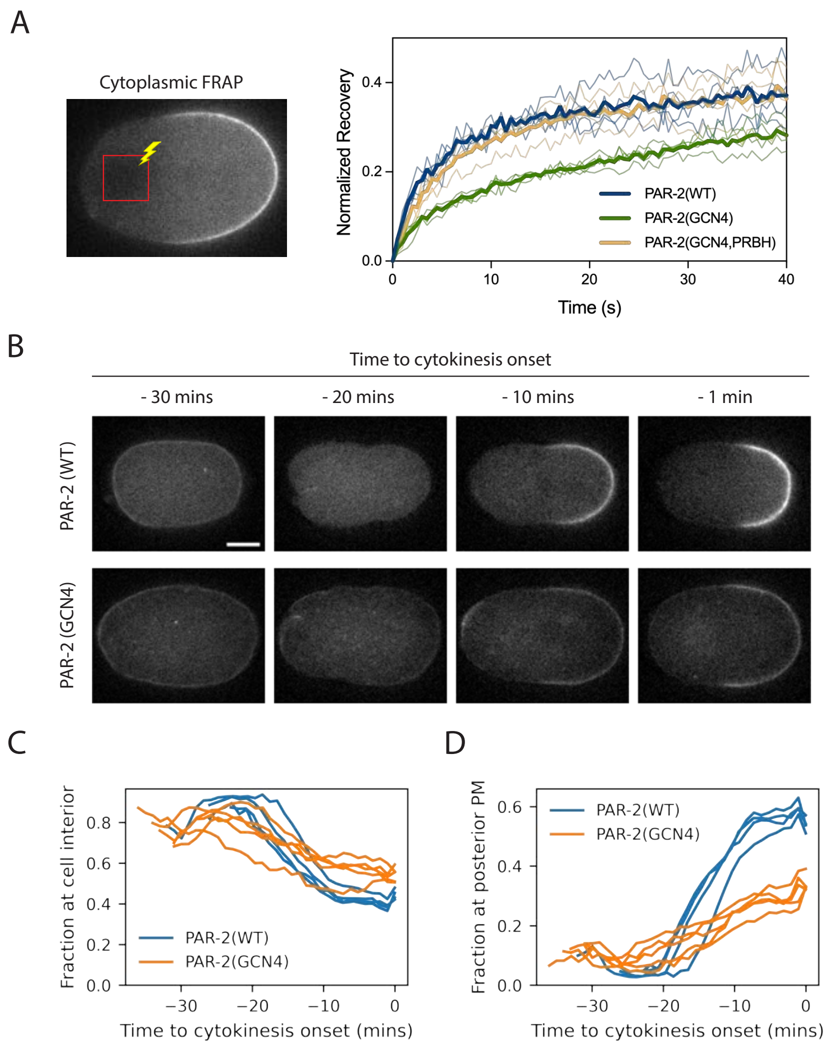

**Figure S8. PAR-2(GCN4) displays reduced mobility compared to wild type. (A)** Normalized recovery curves for cytoplasmic FRAP (see methods). Note that recovery kinetics are reduced for PAR-2(GCN4) compared to wild type, and kinetics are restored by mutating residues in the PRBH site (PRBH). **(B)** - **(D)** PAR-2 localization from meiosis to cytokinesis onset. Images of PAR-2(WT) vs PAR-2(GCN4) (B), along with quantification of the total fraction of PAR-2 at the cell interior (C) and posterior plasma membrane (D), showing a redistribution from the cell interior to the posterior plasma membrane which is slowed for PAR-2(GCN4) compared to wild type. Scale bar in (B) = $10\,\mu m$.

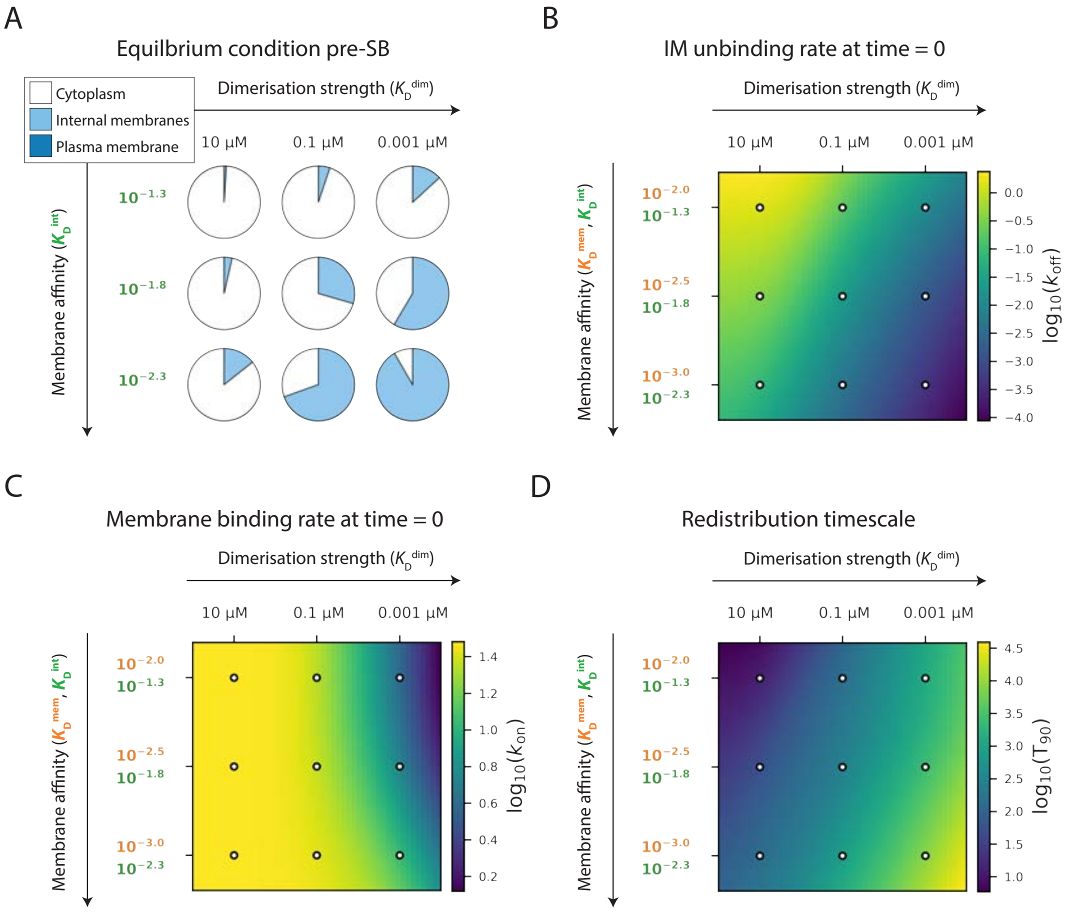

**Figure S9. Entrapment of PAR-2 on internal membranes occurs through dimerization-dependent reduction in membrane exchange rates. (A)** Equilibrium partitioning in the pre-symmetry breaking (SB) equilibrium state, in which PAR-2 is excluded from the plasma membrane. **(B) - (D)** Starting from a pre-SB equilibrium state, $K_D^{mem}$ is decreased at time=0 to simulate the onset of posterior plasma membrane availability, and systems are followed as they progress towards a new equilibrium state. **(B)** Internal membrane (IM) unbinding rate ($k_{off,n}$) at time = 0. White points correspond to the simulations in (A). **(C)** Membrane binding rate ($k_{on}$) at time = 0. **(D)** Redistribution timescale (T90), calculated as the time (in seconds) for plasma membrane concentrations to reach 90% of their post-SB equilibrium level. Note the slowed redistribution kinetics for systems with strong dimerization (low $K_D^{dim}$) as a result of reduced membrane exchange rates.

### Supplemental Tables

**Table S1. Strains and Reagents**

| Name | Description | Source |
|---|---|---|
| Bacteria | | |
| OP50 | E. coli B, ura- | CGC |
| HT115(DE3) | F-, mcrA, mcrB, IN(rrnD-rrnE)1, rnc14::Tn10(DE3 lysogen:lavUV5 promoter-T7 polymerase) | CGC |
| Rosetta(DE3) | F- ompT hsdSB(rB- mB-) gal dcm (DE3) pRARE (CamR) | Novagen |
| Worms | | |
| N2 | Wild type | CGC |
| CGC32 | sC1(s2023) [dpy-1(s2170) umnIs21] III]. | CGC |
| JK2533 | qC1 dpy-19(e1259) glp-1(q339)[qIs26] III/eT1 (III;V) | CGC |
| KK571 | lon-1(e185) par-3(it71)/qC1 dpy-19(e1259) glp-1(q339)III | Cheng et al., 1995 |
| LP637 | par-2(cp329[mNG-C1^PAR-2]) III | Dickinson et al., 2017 |
| NWG0201 | par-2(cp329[mNG-C1^PAR-2]); lon-1(e185) par-3(it71) / qC1 dpy-19(e1259) glp-1(q339)[qIs26] III | This study |
| NWG0240 | par-2(crk41[mNG::par-2(C56S)]*cp329) / sC1(s2023) [dpy-1(s2170) umnIs21] III] | This study |
| NWG0246 | par-2(crk41[mNG::par-2(C56S)]*cp329); lon-1(e185) par-3(it71) / qC1 dpy-19(e1259) glp-1(q339)[qIs26] III | This study |
| NWG0313 | crkSi4[pTB005: mex-5p::GFP::PAR-2(1-177)::PH::nmy-2UTR] | This study |
| NWG0325 | par-2(cp329[mNG-C1^PAR-2]) / sC1(s2023) [dpy-1(s2170) umnIs21] III] | This study |
| NWG0338 | par-2(crk82[mNG::par-2(L109R)]*cp329) / sC1(s2023) [dpy-1(s2170) umnIs21] III] | This study |
| NWG0347 | glh-1(crk148[glh-1::T2A::mNG::INPP4A]) I | This study |
| NWG0351 | par-2(crk89[mNG::par-2(L50R)]*crk82) / sC1(s2023) [dpy-1(s2170) umnIs21] III] | This study |
| NWG0369 | par-2(crk82[mNG::par-2(L109R)]*cp329) lon-1(e185) par-3(it71) / qC1 dpy-19(e1259) glp-1(q339)[qIs26] III | This study |
| NWG0373 | crkSi5[pTB006: mex-5p::GFP::PAR-2(1-177, C56S)::PH::nmy-2UTR] | This study |
| NWG0376 | par-2(crk106[mNG::par-2(GCN4_IL)]*cp329) / sC1(s2023) [dpy-1(s2170) umnIs21] III] | This study |
| NWG0378 | glh-1(crk150[glh-1::tPT2A::mNG::INPP4A]) I | This study |
| NWG0383 | glh-1(crk151[glh-1::tPT2A::mNG::GCN4(IL)]) I | This study |
| NWG0400 | par-2(crk114[mNG::par-2(L50R)]*cp329) / sC1(s2023) [dpy-1(s2170) umnIs21] III] | This study |
| NWG0407 | par-2(crk120[mNG::par-2(C56S, L109R)]*cp329) / sC1(s2023) [dpy-1(s2170) umnIs21] III] | This study |
| NWG0421 | glh-1(crk153[glh-1::tPT2A::mNG::par-2(1-177)]) I | This study |
| NWG0437 | par-2(crk130[mNG::::par-2(GCN4_IL)]*cp329) lon-1(e185)par-3(it71)/qC1dpy-19(e1259)glp-1(q339)[qIs26] III | This study |
| NWG0481 | par-2(crk106[mNG::par-2(6HNL)]*cp329) | This study |
| NWG0489 | par-2(crk170[mNG::par-2(GCN4_IL, S334E, S338E)]*crk106) / sC1(s2023) [dpy-1(s2170) umnIs21] III] | This study |
| NWG0495 | par-2(crk171[mNG::par-2(S334E, S338E)]*cp329) / sC1(s2023) [dpy-1(s2170) umnIs21] III] | This study |
| OD58 | unc-119(ed3) III; ltIs38[pAA1; pie-1::GFP::PH(PLC1 d1) + unc-119(+)] | Audhya et al., 2005 |
| SV2061 | ttTi5605(he314[Ppie-1::glo-epdz::mcherry(smu-1)::tbb-2(3'UTR)]) II; cxTi10816(he259[Peft-3::ph::co-egfp::co- lov::tbb-2(3'UTR)]) IV | Fielmich et al., 2018 |
| Recombinant DNA - RNAi clones | | |
| RNAi Feeding clone: xfp | | C. Eckmann |
| RNAi Feeding clone: par-2 | WB Clone: sjj_F58B6.3 | Source BioScience |
| RNAi Feeding clone: par-6 | WB Clone: sjj_T26E3.3 | Source BioScience |
| RNAi Feeding clone: nop-1 | WB Clone: sjj_F25B5.2 | Source BioScience |
| RNAi Feeding clone: mlc-4 | | Redemann et al, 2010 |
| Recombinant nucleotides - CRISPR | | |
| GFP with introns (for GFP::RING::PH) | ATGAGTAAAGGAGAAGAATTGTTCACTGGAGTTGTCCCAATCCTCGTCGAGCTCGACGGAGACGTCAACGG<br>ACACAAGTTCTCCGTCTCCGGAGAGGGAGAGGGAGACGCCACCTACGGAAAGCTCACCCTCAAGTTCATCTG<br>CACCACCGGAAAGCTCCCAGTCCCATGGCCAACCCTCGTCACCACCTTCTGCTACGGAGTCCAATGCTTCTCCC<br>GTTACCCAGACCACATGAAGCGTCACGACTTCTTCAAGTCCGCCATGCCAGAGGGATACGTCCAAGAGCGTA<br>CCATCTTCTTCAAGGTAAGTTTAAACATATATATACTAACTACTGATTATTTAAATTTTCAGGACGACGGAAAC<br>TACAAGACCCGTGCCGAGGTCAAGTTCGAGGGAGACACCCTCGTCAACCGTATCGAGCTCAAGGTAAGTTT<br>AAACAGTTCGGTACTAACTAACCATACATATTTAAATTTTCAGGGAATCGACTTCAAGGAGGACGGAAACAT<br>CCTCGGACACAAGCTCGAGTACAACTACAACTCCCACAACGTCTACATCATGGCCGACAAGCAAAAGAACGG<br>AATCAAGGTCAACTTCAAGGTAAGTTTAAACATGATTTTACTAACTAACTAATCTGATTTAAATTTTCAGATCC<br>GTCACAACATCGAGGACGGATCCGTCCAACTCGCCGACCACTACCAACAAAACACCCCAATCGGAGACGGAC<br>CAGTCCTCCTCCCAGACAACCACTACCTCTCCACCCAATCCGCCCTCTCCAAGGACCCAAACGAGAAGCGTGAC<br>CACATGGTCCTCCTCGAGTTCGTCACCGCCGCCGGAATCACCCACGGAATGGACGAGCTCTACAAG | |
| PAR-2 1-177 with linkers and introns (for GFP::RING::PH) | GCTAGCATGACCGACCTTGATACCTCCTCCGCCCAAATCAGGGCCAGCCAATCAGACCTATCCATCGCTAACA<br>GCTCCCCACGTTTACTTGAGTCCAACGCGACCACATCCACGGCTTGCATCAACGATGTCCAACAACCTGTCCGT<br>AGAGACCTTTCCAGCGAGCTTCTTTGCCCACTTTGTGACCAACTCTTCGACCGGCCAGTTATGGTGACTTGTG<br>GACACAGCTACTGCGAACCTTGCATCGAGCGCCACACGCGTGACACACGTGCTTGCGTGATCTGCAAGGTAA<br>GTTTAAACATATATATACTAACTAACCCTGATTATTTAAATTTTCAGCTTGATGTCGGACCATTTGCTGCTATG<br>ATCCCATCAATCACGCTCGACAACATGGTTCGCAAGGTAAGTTTAAACAGTTCGGTACTAACTAACCATACAT<br>ATTTAAATTTTCAGCTCAAGAACCAAGAAAACATCGAATCCTCATACGATGACCTGTTCATCTGCGAAGAGAA<br>GCTCCAAAAGGCACCAACTCCACCAATATCTGAAGAGATCCCAGCCCAAAAGATGACTCGCTCTTTATCTCAAC<br>GTTCCAAGGTAAGTTTAAACATGATTTTACTAACTAACTAATCTGATTTAAATTTTCAGAGGCGTCTATTCTTC<br>TCCATCGCTGGGCCAACCATCTGCTCCTCAGCCGGC | |

| | |
|---|---|
| PH (for GFP::RING::PH) | ATGGATTCAGGACGTGATTTCCTTACCCTTCATGGACTCCAAGATGATCCAGACCTTCAAGCTCTTCTTAAAG<br>GATCCCAACTTCTTAAAGTTAAATCATCCTCATGGAGAAGAGAAAGATTCTACAAACTTCAAGAGGATTGTA<br>AGACAATTTGGCAAGAATCAAGAAAGGTTATGAGATCCCCAGAATCACAACTTTTCTCCATTGAAGACATTC<br>AAGAAGTCAGAATGGGACATCGTACAGAAGGACTCGAAAAATTCGCTAGAGATATCCCAGAAGATCGTTGT<br>TTCTCAATCGTCTTCAAAGATCAAAGAAATACACTCGATCTTATTGCTCCATCACCAGCTGACGCTCAACATTG<br>GGTCCAAGGACTTAGAAAGATTATTCATCATTCCGGATCAATGGACCAAAGACAAAAACTTCAACATTGGAT<br>TCACTCATGTCTCAGGAAGGCCGATAAAAACAAAGATAACAAAATGAATTTCAAAGAACTTAAAGACTTCCT<br>TAAAGAACTTAATATCCAATAA |
| mNG with linker and introns (for glh-1 site insertion) | GGAGCATCGGGAGCCTCAGGAGCATCGATGGTCTCCAAGGGAGAGGAGGACAACATGGCCTCCCTCCCAG<br>CCACCCACGAGCTCCACATCTTCGGATCCATCAACGGAGTCGACTTCGACATGGTCGGACAAGGAACCGGAA<br>ACCCAAACGACGGATACGAGGAGCTCAACCTCAAGTCCACCAAGGTAAGTTTAAACATATATATACTAACTA<br>ACCCTGATTATTTAAATTTTCAGGGAGACCTCCAATTCTCCCCATGGATCCTCGTCCCACACATCGGATACGGA<br>TTCCACCAATACCTCCCATACCCAGACGGAATGTCCCCATTCCAAGCCGCCATGGTCGACGGATCCGGATACCA<br>AGTCCACCGTACCATGCAATTCGAGGACGGAGCCTCCCTCACCGTCAACTACCGTTACACCTACGAGGGATCC<br>CACATCAAGGTAAGTTTAAACAGTTCGGTACTAACTAACCATACATATTTAAATTTTCAGGGAGAGGCCCAA<br>GTCAAGGGAACCGGATTCCCAGCCGACGGACCAGTCATGACCAACTCCCTCACCGCCGCCGACTGGTGCCGT<br>TCCAAGAAGACCTACCCAAACGACAAGGTAAGTTTAAACATGATTTTACTAACTAACTAATCTGATTTAAATT<br>TTCAGACCATCATCTCCACCTTCAAGTGGTCCTACACCACCGGAAACGGAAAGCGTTACCGTTCCACCGCCCGT<br>ACCACCTACACCTTCGCCAAGCCAATGGCCGCCAACTACCTCAAGAACCAACCAATGTACGTCTTCCGTAAGAC<br>CGAGCTCAAGCACTCCAAGACCGAGCTCAACTTCAAGGAGTGGCAAAAGGCCTTCACCGACGTCATGGGAA<br>TGGACGAGCTCTACAAG |
| T2A with linker (for glh-1 site insertion) | GGATCGGGAGAGGGACGTGGATCCCTTCTTACCTGCGGAGACGTCGAGGAGAACCCAGGACCA |
| P2A with linker (for glh-1 site insertion) | GGTAGTGGTGCTACCAACTTTTCCCTCTTAAAGCAAGCCGGAGACGTTGAGGAGAATCCAGGACCT |
| INPP4A (for glh-1 site insertion) | TGTATCAGTTCGATATCTGACGG |
| PAR-2 1-177 with linker and introns (for glh-1 insertion) | GGTGGCAGCGGAGGTACCGGCGGTAGTGGAGGCACGATGACCGACCTTGATACCTCCTCCGCCCAAATCAG<br>GGCCAGCCAATCAGACCTATCCATCGCTAACAGCTCCCCACGTTTACTTGAGTCCAACGCGACCACATCCACGG<br>CTTGCATCAACGATGTCCAACAACCTGTCCGTAGAGACCTTTCCAGCGAGCTTCTTTGCCCACTTTGTGACCAA<br>CTCTTCGACCGGCCAGTTATGGTGACTTGTGGACACAGCTACTGCGAACCTTGCATCGAGCGCCACACGCGT<br>GACACACGTGCTTGCGTGATCTGCAAGGTAAGTTTAAACATATATATACTAACTAACCCTGATTATTTAAATT<br>TTCAGCTTGATGTCGGACCATTTGCTGCTATGATCCCATCAATCACGCTCGACAACATGGTTCGCAAGGTAAG<br>TTTAAACAGTTCGGTACTAACTAACCATACATATTTAAATTTTCAGCTCAAGAACCAAGAAAACATCGAATCCT<br>CATACGATGACCTGTTCATCTGCGAAGAGAAGCTCCAAAAGGCACCAACTCCACCAATATCTGAAGAGATCCC<br>AGCCCAAAAGATGACTCGCTCTTTATCTCAACGTTCCAAGGTAAGTTTAAACATGATTTTACTAACTAACTAAT<br>CTGATTTAAATTTTCAGAGGCGTCTATTCTTCTCCATCGCTGGGCCAACCATCTGCTCCTCA |
| GCN4 with linkers (for par-2 and glh-1 site insertion) | GGATCCGGAGGACGTATGAAACAGCTTGAGGACAAGATTGAGGAATTACTCTCCAAGATTTATCACCTCGA<br>GAACGAGATTGCCCGTCTCAAAAAATTAATCGGTGAGCGTGGTGGATCCGGT |
| 6HNL with homology arms, linkers and introns (for par-2 insertion) | TTAGCTCAAAAACCCAATTTTCTAGCTAAAATCTCAAATTTTTGCATTTTTTCGCAGAAAAATACCAAAACATTC<br>AACGAATTTTCGTTTCAGATGGTCCGAAAATTGAAAAACCAAGAGGGCAGCGGAGGTACCGGCGGTAGTG<br>GAGGCGGCAGTCATATGGCTCATCAACCATCCGACACCATCGCCGGACTCTACGAAGCCTTCAATTCGGGAG<br>ATTTGGAGACTCTTCGTGAGCTCATCGCTCCAGATGCCGTCATCCACCTTCCCGGCACCGCCGGAGGTAAGTT<br>TTGCCGGGATTATTCATATACATTAATAATAATGGCTTTTTCAGATGCTGAGCACCCACCAGGAACTCCACGC<br>GATCGTGAAGGATGGTTGGGAGTTTGGCAGTTCACCCAGGCTTTCTTCCCCGATATGACAGCGACCGTTCA<br>GGTAAGTTTTGATCGGGAAGTGATAAATCGTAAGTTGAAAAAAAATTTTCAGGACATCGTCCAAACCGGTG<br>ATCTTGTTGCTACTCGTTGTGTGGCTCGCGGAACACACAGTGGACGTCCGTTCGAGATGACCATGTTGAACA<br>TGAGTCGTGTTAGAGATGGACGCATTGTCGAGCACTGGACGATCAGCGACAACGTCACCATGCTTGCTCAA<br>CTTGGAGTCAAGGCCTCACTCGGTGGATCCGGTAATATCGAGAGCAGCTACGACGATTTGTTCATTTGTGA<br>GGAAAAGGTTGAGTTTTTGCAGGAAAATGCGAATTTTCAGGGTTTTTGAGTGAAAAAATCATGAAAATTA<br>GGTGTTTT |
| sgRNA for par-2 C56S, L50R | GCTGATCACACAGTGGACAG |
| sgRNA for par-2 C56S, L50R | TCGAAAAGCTGATCACACAG |
| sgRNA for par-2 L109R | GTAATCTGTAAACTCGACGT |
| sgRNA for par-2 L109R | CTGGGAATCATCGCTGCAAA |
| sgRNA for par-2 GCN4, 6HNL insertions | AGCTGCTCTCGATATTCTCC |
| sgRNA for par-2 GCN4, 6HNL insertions | GGTCCGAAAATTGAAAAACC |
| sgRNA for par-2 PRBH mutations | GCGGCTTTTCATGCCGAAAA |
| sgRNA for par-2 PRBH mutations | GGTGTCAAAAGTGTCGCCAG |
| sgRNA for mNG insertion at glh-1 | CAAGTCCCTCAAGATGAAGA |
| sgRNA for mNG insertion at glh-1 | TCCCTCAAGATGAAGAAGGC |
| sgRNA for P2A insertion at glh-1 | GGACGAGGAGGGGTGGGGAT |
| sgRNA for P2A insertion at glh-1 | GGAGGGGTGGGGATCGGGAG |
| sgRNA for INPP4A | TGTATCAGTTCGATATCTGA |
| par-2 C56S repair template | TGCATCAACGACGTTCAACAGCCGGTTCGACGCGATTTGAGCTCGGAACTCTTAAGCCCCCTGTGTGATCAAT<br>TGTTCGACAGGGTTAGAACATGGAAA |
| par-2 L50R repair template | TGCATCAACGACGTTCAACAGCCGGTTCGACGCGATAGGAGCTCGGAACTCTTATGTCCCCTGTGTGATCAA<br>TTGTTCGACAGGGTTAGAACATGGAAA |
| par-2 L109R repair template | GATACGAGAGCCTGTGTTATATGCAAACTGGACGTTGGACCATTCGCTGCCATGATCCCTAGCATTACACGT<br>GATAATGTACGACGGATTTTGATGCGAA |
| par-2 PRBH repair template | AGGAGACGTGGAGCATTTGCTCCCAGAGAAGCCGACCAAAGAAAGCATAAGAGAGCTCTGGAAGCGGCTT<br>TTCATGCCGAAAAAAGTGTCAAAAGTGTCGCCCGCAGCCGTCCAGCTGCTGAGCCAATCGCCCAGCACCACT<br>GACGAACGGAGCCACC |
| par-2 GCN4 repair template | TTTTCGTTTCAGATGGTCCGAAAATTGAAAAACCAAGAGGGATCCGGAGGACGTATGAAACAGCTTGAGG<br>ACAAGATTGAGGAATTACTCTCCAAGATTTATCACCTCGAGAACGAGATTGCCCGTCTCAAAAAATTAATCG<br>GTGAGCGTGGTGGATCCGGTAATATCGAGAGCAGCTACGACGATTTGTTCATTTGTGA |
| glh-1 P2A repair template | CAATTACGGAGCTAGTGGATTTGGGTCCAGTGTACCAACTCAAGTCCCGCAGGACGAGGAGGGGTGGGGT<br>AGTGGTGCTACCAACTTTTCCCTCTTAAAGCAAGCCGGAGACGTTGAGGAGAATCCAGGACCTGGATCGGG<br>AGAGGGACGTGGATCCCTTCTTACCTGCGGAGACGTCGAGGAGAACCCAGGACCAGGAG |

| | | |
|---|---|---|
| glh-1 GCN4 repair template | GCCTTCACCGACGTCATGGGAATGGACGAGCTCTACAAGGGATCCGGAGGACGTATGAAACAGCTTGAGGACAAGATTGAGGAATTACTCTCCAAGATTTATCACCTCGAGAACGAGATTGCCCGTCTCAAAAAATTAATCGGTGAGCGTGGTGGATCCGGTTAGAAAACCGACCAATTGATAGTGTTTCGCATTTATTA | |
| fwd primer for mNG + T2A + INPP4A PCR | GGATCGGGAGAGGGACGTGGATCCCTTCTTACCTGCGGAGACGTCGAGGAGAACCCAGGACCAGGAGCATCGGGAGCCTCAGGAGCATCGATGGTCTCCAAGGGAGAGGA | |
| rev primer for mNG + T2A + INPP4A PCR | TCCGTCAGATATCGAACTGATACACTTGTAGAGCTCGTCCATTC | |
| fwd primer for mNG + T2A + INPP4A + homology PCR | TGTTCCAGACTGGATGCAAGGTGCTGCTGGAGGCAATTACGGAGCTAGTGGATTTGGGTCCAGTGTACCAACTCAAGTCCCGCAGGACGAGGAGGGGTGGGGATCGGGAGAGGGACGTGG | |
| rev primer for mNG + T2A + INPP4A + homology PCR | CACCAACAAATACAATTAAAAATCAACAAGGGCAGGATAAAATATGGGGGAACTGACAGCATTAATAAATGCGAAACACTATCAATTGGTCGGTTTTCTATCCGTCAGATATCGAACTGA | |
| fwd primer for PAR-2 1-177 PCR | GGTGGCAGCGGAGGTACCGGCGGTAGTGGAGGCACGATGACCGACCTTGATACCTC | |
| rev primer for PAR-2 1-177 PCR | TGAGGAGCAGATGGTTGGCC | |
| fwd primer for PAR-2 1-177 + homology PCR | CTTCCGTAAGACCGAGCTCAAGCACTCCAAGACCGAGCTCAACTTCAAGGAGTGGCAAAAGGCCTTCACCGACGTCATGGGAATGGACGAGCTCTACAAGGGTGGCAGCGGAGGTACCGGCGGTAGTGGAGGCACGATGACCGACCTTGATACCTC | |
| rev primer for PAR-2 1-177 + homology PCR | CACCAACAAATACAATTAAAAATCAACAAGGGCAGGATAAAATATGGGGGAACTGACAGCATTAATAAATGCGAAACACTATCAATTGGTCGGTTTTCTATGAGGAGCAGATGGTTGGCC | |
| fwd primer for 6HNL PCR | GGCAGCGGAGGTACCGGCGG | |
| rev primer for 6HNL PCR | ACCGGATCCACCGAGTGAGG | |
| fwd primer for 6HNL + homology PCR | TTAGCTCAAAAACCCAATTTTCTAG | |
| rev primer for 6HNL + homology PCR | AAAACACCTAATTTTCATGATTTTTTCA | |
| fwd primer for par-2 1-177 C56S site directed mutagenesis | CGAGCTTCTTAGCCCACTTTG | |
| rev primer for par-2 1-177 C56S site directed mutagenesis | CTGGAAAGGTCTCTACGG | |
| fwd screening primer for par-2 C56S, L50R | GATTGCCAACTCATCGCCAC | |
| rev screening primer for par-2 C56S, L50R | TCCGGCAAAATTGGGGTTTT | |
| fwd screening primer for par-2 L109R | AGAAACCCCAGTTTTTCAGCG | |
| rev screening primer for par-2 L109R | TGAATTTTCGGCAAGATTTTCAAGGA | |
| fwd screening primer for par-2 PRBH mutations | CGCTCTCACCTCCAGGATTC | |
| rev screening primer for par-2 PRBH mutations | ATTTTAGAACGAACGGCGGC | |
| fwd screeing primer for par-2 GCN4, 6HNL insertions | CATGTGGGCACTCGTACTGT | |
| rev screeing primer for par-2 GCN4, 6HNL insertions | AAACCCGACTTTTGGGGTCA | |
| fwd screening primer for glh-1 insertions (mNG, P2A) | TCGCTGAACGTGGACTTGAT | |
| fwd screeing primer for glh-1 insertions (RING, GCN4) | CTACACCACCGGAAACGGAA | |
| rev screening primer for glh-1 insertions (mNG, RING, GCN4) | CCGCGGAGAATCGGAAAACA | |
| rev screeing primer for glh-1 insertions (P2A) | GCCATGTTGTCCTCCTCTCC | |
| Recombinant DNA - Protein Biochemistry | | |
| par-2 40-120 | AACGACGTTCAACAGCCGGTTCGACGCGATTTGAGCTCGGAACTCCTCTGTCCACTGTGTGATCAGCTTTTCGACAGGCCCGTAATGGTAACATGTGGGCACTCGTACTGTGAGCCGTGCATCGAACGACATACACGTGATACGAGAGCCTGTGTAATCTGTAAACTCGACGTTGGACCATTTGCAGCGATGATTCCCAGCATTACACTTGATAATATGGTCCGAAAATTGAAAAACCAGGAG | |
| fwd primer for par-2 40-120 L109R site directed mutagenesis | CAGCATTACACGCGATAATATGGTCCG | |
| rev primer for par-2 40-120 L109R site directed mutagenesis | GGAATCATCGCTGCAAATG | |
| Plasmids | | |
| pRI021 | ttTi5605 Mos1 insertion vector (mex-5 promoter and nmy-2 3' UTR) | This study |
| pDD122 | Cas9 + sgRNA plasmid for ttTi5605 Mos1 insertion | Dickinson et al., 2013 |
| pETM11-SUMO3eGFP | His-SUMO bacterial expression vector | EMBL |

| Genotype | Normalised cytoplasmic concentration (μm$^{-3}$) | Normalised posterior membrane concentration (μm$^{-2}$) | Posterior M:C ratio (μm) | Fraction at membrane | Total expression (μm$^{-3}$) |
|---|---|---|---|---|---|
| PAR-2(WT) | 1.00 ± 0.13 | 22.91 ± 3.80 | 22.90 ± 2.46 | 0.62 ± 0.03 | 2.62 ± 0.37 |
| PAR-2(C56S) | 1.73 ± 0.14 | 5.46 ± 1.25 | 3.17 ± 0.72 | 0.16 ± 0.03 | 2.06 ± 0.17 |
| PAR-2(L109R) | 1.62 ± 0.23 | 9.99 ± 1.79 | 6.19 ± 0.71 | 0.28 ± 0.03 | 2.25 ± 0.33 |
| PAR-2(WT); par-3(it71) | 0.92 ± 0.14 | 12.41 ± 1.69 | 13.65 ± 1.78 | 0.66 ± 0.03 | 2.70 ± 0.32 |
| PAR-2(C56S); par-3(it71) | 1.35 ± 0.14 | 5.08 ± 0.61 | 3.80 ± 0.56 | 0.39 ± 0.03 | 2.19 ± 0.19 |
| PAR-2(L109R); par-3(it71) | 1.42 ± 0.21 | 8.63 ± 1.60 | 6.09 ± 0.79 | 0.48 ± 0.02 | 2.73 ± 0.40 |

**Table S2. Table of quantification results for PAR-2(WT), PAR-2(C56S) and PAR-2(L109R) in polarized and uniform (par-3(it71)) conditions.** Mean ± SD.

| ***Hydrodynamic parameters*** | |
|---|---|
| ***Protein*** | PAR-2 RING |
| $^{a}v$ (mL.g$^{-1}$) | 0.728 |
| $^{b}\rho$ (g.mL$^{-1}$) | 1.005 |
| $^{c}\eta$ (x10$^{2}$) (g$^{-1}$ cm$^{-1}$ s$^{-1}$) | 1.022 |
| $^{d}M_r$ | 9,235 |
| $^{e}\varepsilon_{280}$ (M$^{-1}$ cm$^{-1}$) | 1,300 |
| $^{f}J_{inc}$ (M$^{-1}$.cm$^{-1}$) | 25,396 |

[a]Protein partial specific volume; [b]Buffer density; [c]Buffer viscosity [d]Molar mass calculated from the protein sequence; [e]Molar absorbance extinction coefficient; [f]Molar fringe increment.

| ***Sedimentation equilibrium*** | | | | |
|---|---|---|---|---|
| ***PAR-2 RING*** | | | | |
| C (μM) | 85 | 125 | 240 | 85-240 |
| $^{a}M_w$ kD | 19.2 | 18.8 | 18.0 | 18-19.2 |
| $^{b}K_D^{dim}$ (μM) | 0.36 | 0.37 | 0.33 | 0.36 |
| $^{c}$rmsd | 0.006 | 0.006 | 0.004 | 0.004 – 0.006 |
| $^{d}\chi^2$ | | | | 1.25 |

[a]weight averaged molecular weight derived from Global analysis of individual samples using single species model; [b]monomer-dimer equilibrium dissociation constant determined from a global fit using three concentrations and two speeds to a monomer-dimer self-association model; [c]rms deviation observed for each multi-speed sample when fitted individually and globally; [d]global reduced chi-squared from combined fitting of all multispeed data.

**Table S3. PAR-2 RING sedimentation equilibrium data.**

| a (nm) | $K_D^{dim}$ WT fixed? | $K_D^{dim}$ WT (μM) | $K_D^{dim}$ L109R (μM) | $K_D^{dim}$ fold difference | $\log_{10}(K_D^{mem})$ | Figure |
|---|---|---|---|---|---|---|
| 5 | x | 0.425 [0.275, 0.712] | 2.460 [1.547, 4.415] | 5.79 [5.26, 6.53] | -2.43 [-2.52, -2.34] | 3F |
| 0.5 | x | 3.580 [1.633, 6.602] | 19.423 [8.026, 40.807] | 5.43 [4.86, 6.21] | -3.38 [-3.51, -3.22] | S6B |
| 50 | x | 0.095 [0.082, 0.114] | 0.750 [0.632, 0.908] | 7.90 [7.05, 8.93] | -1.62 [-1.65, -1.60] | S6C |
| 5 | ✓ | 0.358 | 2.043 [1.866, 2.231] | 5.71 [5.21, 6.23] | -2.39 [-2.41, -2.38] | S6D |
| 0.5 | ✓ | 0.358 | 1.703 [1.562, 1.850] | 4.76 [4.36, 5.17] | -2.91 [-2.92, -2.90] | S6E |
| 50 | ✓ | 0.358 | 3.902 [3.263, 4.915] | 10.90 [9.11, 13.73] | -1.83 [-1.84, -1.82] | S6F |

**Table S4. Thermodynamic model parameters.** Optimized parameters from six different fits of the thermodynamic model to the in vivo PAR-2(WT) and PAR-2(L109R) rundown data. Shows results with/without fixing the wild type $K_D^{dim}$ to the value experimentally determined by AUC, and with three different values for the protein diameter *a*. 95% confidence intervals determined by bootstrapping.
