## Supplemental Methods for "Optimized dimerization of the PAR-2 RING domain drives cooperative and selective membrane recruitment for robust feedback-driven cell polarization"

#### Materials and Methods

##### C. elegans - strains and culture conditions

*C. elegans* strains were maintained at 20°C on nematode growth media (NGM) seeded with OP50 bacteria (Stiernagle, 2006). Strains are listed in Table S1.

##### Strain construction

To generate point mutations or small insertions, mutation by CRISPR/Cas9 was performed using the protocol described by Arribere et al. (2014). Repair templates were designed containing the target mutation and silent restriction sites to aid screening. crRNA guides were annealed with tracrRNA (IDT) by combining 0.5 μl tracrRNA (4 mg/ml) with 2.75 μl guide (100 μM) and 2.75 μl duplex buffer (IDT), and incubating at 95°C for 5 minutes. Injection mixes were prepared containing the annealed crRNA/tracrRNA, repair template and Cas9 protein (IDT, 1 μL at 10 mg/ml), along with either *dpy-10* or *unc-58* co-CRISPR markers (Arribere et al., 2014). Injection mixes were incubated at 37°C for 10 minutes, centrifuged at 14,000 rpm for 10 minutes and injected into the gonads of adult worms. Mutants were identified by PCR and verified by sequencing.

To generate the membrane-tethered RING domain construct, sequences for PH, GFP and PAR-2(1-177) were assembled in pRI21, a vector designed for inserting genes at the ttTi5605 *mos1* locus under control of a *mex-5* promoter and an *nmy-2* 3' UTR. A C56S mutant form of the construct was generated by site directed mutagenesis (Q5 site directed mutagenesis kit from New England Biolabs). Insertions were performed by CRISPR using the method described by Dickinson et al. (2015).

mNG, mNG::RING and mNG::GCN4 were expressed by inserting at the 3' end of the *glh-1* gene preceded by a self-cleaving peptide using an approach similar to Goudeau et al. (2021). NeonGreen was inserted first, flanked by the self-cleaving peptide T2A and INPP4A, an optimized CRISPR guide site included to serve as a base for further insertions (Duan et al., 2020). Insertion was performed by CRISPR/Cas9 using the method described by Dokshin et al. (2018). crRNA guides targeting the 3' end of *glh-1* were annealed with tracrRNA (IDT) by combining 0.5 μl tracrRNA (4 mg/ml) with 2.75 μl guide (100 μM) and 2.75 μl duplex buffer (IDT), and incubating at 95°C for 5 minutes. PCR products containing the sequence to be inserted (tPT2A::mNG::INPP4A), and the same sequence with 100bp homology arms to *glh-1* were generated and column purified (Qiagen, QIAquick PCR purification kit), mixed in equimolar amounts, denatured at 95°C, and annealed by gradually cooling to room temperature to generate a pool of products with single-stranded DNA overhangs to act as the repair template. The injection mix was prepared containing the annealed crRNA/tracrRNA, repair template and Cas9 protein (IDT, 1 μL at 10 mg/ml), along with a *dpy-10* co-CRISPR marker (Arribere et al., 2014). The injection mix was incubated at 37°C for 10 minutes, centrifuged at 14,000 rpm for 10 minutes and injected into the gonads of adult worms. Mutants were identified by PCR and verified by sequencing. Whilst this led to good expression of mNG, we found considerable expression of GLH-1::mNG fusion products resulting from incomplete ribosome skipping at T2A (Kim et al., 2011). To minimize this, we inserted an additional self-cleaving peptide, P2A, in tandem with T2A (Liu et al., 2017; Pan et al., 2017) by CRISPR/Cas9, which reduced the incidence of read-through products compared to T2A alone, without impacting expression levels. Further insertions (RING and GCN4) were performed by CRISPR/Cas9 targeted to the INPP4A site, generating N-terminal mNG fusions (mNG::RING and mNG::GCN4). Recombinant oligonucleotides are listed in Table S1.

##### RNA interference

RNAi was performed using the feeding method described in Kamath et al. (2003). Bacterial feeding clones were grown in LB liquid culture with ampicillin (50 μg/ml) for 16 hours at 37°C in a shaking incubator. dsRNA expression was then induced with IPTG (5 mM), and 150 ml bacteria were struck onto 60 mm NGM agar plates, which were then incubated at room temperature for 24 hours. To obtain complete depletions, L4 worms were added to plates and incubated at 20°C for 24-48 hours before imaging. To obtain graded depletions, adult/L4 worms were left on plates for 0-24 hours.

##### Microscopy

Embryos were dissected in 8 μl egg buffer or Shelton's Growth Medium for meiotic embryos (Shelton and Bowerman, 1996), and mounted between a slide and coverslip with 20 μm polystyrene beads. Midplane confocal images were captured using a 60x objective lense on a Nikon TiE system equipped with an X-Light V1 spinning disk system (CrestOptics) with 50 um slits, Obis 488/561 nm fiber-coupled diode lasers (Coherent) and an Evolve delta camera (Photometrics). The system was controlled using Metamorph (Molecular Devices) and configured by Cairn Research. For photobleaching, embryos were mounted as above, but imaged using a 100x 1.4 NA objective. Photobleaching was performed using a 473 nm laser directed by an iLAS targeted illumination system (Roper). A 50 x 50 px box was bleached in the center of the anterior cytoplasm and images captured at 0.5s intervals.

##### Image analysis - FRAP

Fluorescence within the bleached ROI and a corresponding control ROI in the posterior cytoplasm were measured. Fluorescence in the bleached ROI was first normalized to the control ROI and then normalized to the prebleach and postbleach signals. Prebleach was defined by the mean signal of the 10 frames prior to bleaching. As all embryos experienced a very rapid initial recovery phase, we defined the postbleach frame as 1 s after bleaching to isolate long timescale kinetics.

##### Image analysis - quantification of membrane and cytoplasmic concentrations

*Image preprocessing*

Images were autofluorescence corrected using SAIBR (Rodrigues et al., 2022), and a 50-pixel wide line following the membrane around the embryo was computationally straightened to simplify geometry for further analysis.

*Quantification model*

Our method is adapted from previous methods that model intensity profiles perpendicular to the membrane as the sum of distinct cytoplasmic and membrane signal components (Gross et al., 2018; Reich et al., 2019). Typically these two components are modelled as an error function and Gaussian function respectively, representing the expected shape of a step and a point convolved by a Gaussian point spread function (PSF) in one dimension. Using this model, one can generate simulated images of straightened cortices as the sum of two tensor products which represent distinct membrane and cytoplasmic signal contributions (Figure S1A):

$$c_{\text{cyt}} \otimes s_{\text{cyt}} + c_{\text{mem}} \otimes s_{\text{mem}}$$

where $c_{cyt}$ and $c_{\text{mem}}$ are cytoplasmic and membrane concentration profiles and $s_{\text{mem}}$ and $s_{\text{cyt}}$ are, by default, Gaussian and error function profiles. We impose the constraint that the cytoplasmic concentration $c_{\text{cyt}}$ is uniform throughout each image. Using a differentiable programming paradigm, the input parameters to the model can be iteratively adjusted by backpropagation to minimize the mean squared error between simulated images and ground truth images. As well as allowing the image-specific concentration parameters ($c_{cyt}$ and $c_{\text{mem}}$) to be learnt, this procedure also allows the global signal profiles $s_{\text{mem}}$ and $s_{\text{cyt}}$ to be optimised and take any arbitrary form, allowing the model to generalise beyond a simple Gaussian PSF model and account for complex sample-specific light-scattering behaviors. We describe this procedure below. In practice we found that this additional flexibility is necessary to minimise model bias and prevent underfitting (Figure S1F).

Analysis was performed in Python using the differentiable programming package JAX (Bradbury et al., 2018). All optimizations were performed with an Adam optimizer (Kingma and Ba, 2015) and a learning rate of 0.005, and run until the loss function (mean squared error) was stabilized.

*Model training*

Model training was performed in a two step process (Figure S1B). $s_{\text{cyt}}$ was trained by performing gradient descent on images of cytoplasmic NeonGreen protein with the $s_{\text{cyt}}$ and $c_{cyt}$ as free parameters, and the membrane signal contribution fixed to zero. Training was performed on 17 images in batch with a shared $s_{\text{cyt}}$, resulting in the optimized $s_{\text{cyt}}$ profile shown in Figure S1C.

$s_{\text{mem}}$ was trained by performing gradient descent on images of polarized PAR-2 with $s_{\text{mem}}$, $c_{\text{mem}}$ and $c_{cyt}$ as free parameters, and $s_{\text{cyt}}$ fixed to the previously determined profile. The use of polarized images, along with the assumption of a uniform cytoplasmic contribution, allows the model to learn $s_{\text{mem}}$ based on the difference in signal between the anterior and posterior halves of the embryo. To test the generalizability of the model, we performed training on separate batches of wild type PAR-2, PAR-2(L109R) and PAR-2(C56S) images, as well as heterozygous PAR-2 images with a single mNG-tagged copy (50% signal). Training was performed on 10 images with a shared $s_{\text{mem}}$ for each batch. We found the resulting shape to be similar between all lines (Figure S1D). Notably, profiles are asymmetric, with a higher signal at the internal portion of the curve. We reason that this is due to out of focus contributions from membrane protein above and below the plane of the image (Figure S1E), which is not accounted for by previous methods. For subsequent quantification steps we used a model trained on the full dataset of 50 images (10 for each condition) (Figure S1D, black line).

*Quantification*

With $s_{\text{cyt}}$ and $s_{\text{mem}}$ fixed to the values determined above, quantification was performed on images by performing gradient descent with $s_{\text{cyt}}$ and $s_{\text{mem}}$ fixed and $c_{cyt}$ (the uniform cytoplasmic concentration) and $c_{\text{mem}}$ (an array of membrane concentrations around the embryo) optimized as free parameters.

*Calibration of membrane and cytoplasmic units*

$c_{cyt}$ and $c_{\text{mem}}$ at this point are in their own arbitrary units, and so a conversion parameter is required to put them into common units. To calibrate this conversion parameter, we quantified the effects on raw membrane and cytoplasm concentration measurements of redistributing a fixed pool of protein from the cytoplasm to the membrane, using an optogenetics PH::eGFP::LOV/ePDZ::mCherry system (Fielmich et al., 2018). Embryos were exposed to blue light for 10 seconds to promote an interaction between ePDZ and LOV, leading to a rapid uniform recruitment of ePDZ::mCherry to the membrane and a rebalancing of the total protein pool (Figure S1G). Expressing the total pool of protein before and after blue light exposure as:

$$T = C + \psi c M \text{ (before)} \qquad T = C' + \psi c M' \text{ (after)}$$

where $C/C'$ and $M/M'$ are mean membrane and cytoplasmic concentrations in raw model units before/after exposure, the unit conversion factor $c$ can be calculated by comparing the gain in $M$ post-exposure to the loss in $C$:

$$c = \frac{C - C'}{\psi(M' - M)}$$

Full quantification data for wild-type, C56S and L109R PAR-2 in *par-3(wt)* and *par-3(it71)* conditions is shown in Table 1. Membrane and cytoplasmic concentrations have been converted to equivalent units using the conversion parameter $c$, and all measures have been normalized to the cytoplasmic concentration of wild type PAR-2 in *par-3(wt)* conditions. Here and throughout the paper, posterior membrane concentrations are defined as the mean concentration within the posterior-most 20% of the plasma membrane. Posterior M:C ratio is defined as the posterior membrane concentration divided by the (uniform) cytoplasmic concentration. Fraction at the PM is defined as the amount of protein at the plasma membrane divided by the total amount in the cell (cytoplasmic + membrane). Fraction at the posterior PM is the corresponding measure limited to the posterior-most 50% of the PM. Peak membrane concentration in Figure 4E is defined as the highest concentration within any 20% area of the membrane.

**Scoring cooperativity**

We consider a system in which protein exchanges between the cytoplasm ($c$) and membrane ($m$) with the following governing equation:

$$\frac{dm}{dt} = k_{\text{on}} c - k_{\text{off}} m$$

where $k_{\text{on}}$ and $k_{\text{off}}$ are membrane binding and unbinding rates. At equilibrium ($\frac{dm}{dt} = 0$), the following condition holds:

$$m = \frac{k_{\text{on}}}{k_{\text{off}}} c$$

We consider a cooperative system in which $k_{\text{on}}$ and/or $k_{\text{off}}$ vary as a function of $m$. The precise form will depend on mechanistic details, but, for the purposes of scoring cooperativity, we assume that the quantity $k_{\text{on}}/k_{\text{off}}$ will take the general form of an exponential ($\beta m^{\lambda}$) across the relevant range of concentrations. We can then rewrite the equilibrium condition as

$$m = \beta c^{\alpha}$$

where $\alpha = 1/(1-\lambda)$. For example, a system in which $k_{\text{on}}$ is proportional square-root of membrane concentration (with constant $k_{\text{off}}$) will have $\lambda = 0.5$, $\alpha = 2$, whereas a linear system will have $\lambda = 0$, $\alpha = 1$. Here, we can see that, for any $\alpha \neq 1$, equilibrium ratios between $m$ and $c$ will be concentration-dependent.

To score the cooperativity of in vivo systems, we used graded RNAi, along with the image quantification procedure described previously, to quantify $m$ and $c$ in systems with varying quantities of protein. Then, using a log-transformed version of the equilibrium condition

$$\log_{10}(m) = \alpha \log_{10}(c) + \beta$$

we quantified $\alpha$ and $\beta$ by performing linear regression on the log-transformed data, with $\alpha$ given as the cooperativity score. Probability distributions for $\alpha$ were calculated by bootstrapping.

**RING domain expression and purification**

The DNA sequence for PAR-2 residues 40-120 (containing the core RING domain and flanking dimerization helices) was amplified from plasmid pNH46 and cloned into the pETM11 His-Sumo vector (provided by the Crick Structural Biology STP). An L109R mutant construct was generated by site directed mutagenesis. Plasmids were verified by sequencing.

Protein was expressed in Rosetta (DE3) cells overnight at 16°C in LB media supplemented with 50 mM zinc sulphate. Protein was then purified by affinity chromatography (Ni-NTA agarose kit from Qiagen), and the tag was removed by treatment with SenP2 protease (provided by the Crick Structural Biology STP). Protein was further purified by ion exchange chromatography (Cytiva Hi-TrapMonoQ 1 ml column) and size exclusion chromatography (Cytiva Superdex 75 increase 10/30 column).

**SEC-MALS**

Samples were run over a size exclusion chromatography using a Superdex 75 increase 10/30 column (Cytiva), and analysed by multi-angle light scattering (MALS) using a DAWN MALS detector (Wyatt instruments). Molecular weight measurements in Figure 2H are shown as a weight-average. For a system containing a mix of monomers and dimers, this can be modelled as:

$$M_w = \frac{n_m W_m^2 + n_d W_d^2}{n_m W_m + n_d W_d}$$

where $W_m$ and $W_d$ are the molecular weight of monomer and dimer molecules (= 9.23474 and 18.46948 kDa for the PAR-2 RING domain), and $n_m$ and $n_d$ (the number of monomer and dimer molecules in the sample) can be described as a function of total concentration using a dimerization model (see supplemental model description).

**Analytical ultracentrifugation**

Sedimentation equilibrium experiments were performed in a Beckman Optima-AUC analytical ultracentrifuge using aluminium double sector centrepieces in an An-50 Ti rotor. Solvent density and the protein partial specific volumes were determined as described (Laue et al., 1992). Prior to centrifugation, PAR-2 RING samples were dialyzed exhaustively against the buffer blank (20 mM Tris-HCl, 150 mM NaCl, 0.5 mM TCEP). Samples (150 µL) and buffer blanks (160 µL) were loaded into the cells and after centrifugation for 30 hours at 20,000 rpm interference data were collected at 2 hourly intervals until no further change in the profiles was observed.

The rotor speed was then increased to 24,000 rpm, and the procedure repeated. Data were collected on samples at three different PAR-2 RING concentrations. The program SEDPHAT (Vistica et al., 2004) was used to initially determine weight-averaged molecular masses by nonlinear fitting of individual multi-speed equilibrium profiles to a single-species ideal solution model. Inspection of these data revealed that the molecular mass showed significant increase over the monomer molecular weight. Therefore, global fitting of the data to a monomer-dimer model incorporating the data from multiple speeds and multiple sample concentrations was applied to extract the monomer-dimer equilibrium dissociation constant ($K_D^{dim}$).

**Structure prediction**

The PAR-2 RING structure was predicted using AlphaFold-Multimer (Jumper et al., 2021; Evans et al., 2022) via the online AlphaFold Colab notebook, using a dimer model with residues 40-120 of PAR-2. High prediction confidence (pLDDT $> 90$) was reported for the majority of the output (residues L50 to K117).

**Modelling - equilibrium model**

*Four species thermodynamic model*

We consider a system in which protein is both dimerizing and exchanging between membrane and cytoplasmic pools. There are four species to consider: membrane monomers ($m_1$), membrane dimers ($m_2$), cytoplasmic monomers ($c_1$) and cytoplasmic dimers ($c_2$). The membrane is modelled as a thin volume compartment with thickness $a$ equal to the protein diameter (Figure S6A), with protein exchanging between this compartment and the underlying cytoplasm. Thus, total protein amounts $c_{tot}$ are conserved according to

$$c_{tot} = c_c + a\psi c_m \tag{1}$$

where $c_c$ $(= c_{c_1} + c_{c_2})$ and $c_m$ $(= c_{m_1} + c_{m_2})$ are protein concentrations in the cytoplasmic and membrane compartments, and $\psi$ is the membrane surface area to cytoplasmic volume ratio. Protein diameter $a$ has not been experimentally determined for PAR-2, but given a molecular weight of 69.95 kDa, the diameter is expected to be approximately 5 nm (Erickson, 2009). Chemical equilibrium is defined by the following condition:

$$\mu_c = \mu_m \tag{2}$$

where $\mu_m$ and $\mu_c$ are effective chemical potentials for the membrane and cytoplasm, which take the following form when we assume dimerization equilibrium (see supplemental model description for full derivation):

$$\frac{\mu_c}{RT} = \ln\left(\frac{c_c}{c_0}\right) - \frac{1}{2}\ln\left(1 + \frac{4c_c}{K_D^{dim}} + \sqrt{1 + \frac{8c_c}{K_D^{dim}}}\right) + 1 + s_0$$
$$\frac{\mu_m}{RT} = \ln\left(\frac{c_m}{c_0}\right) + \ln(K_D^{mem}) - \frac{1}{2}\ln\left(1 + \frac{4c_m}{K_D^{dim}} + \sqrt{1 + \frac{8c_m}{K_D^{dim}}}\right) + 1 + s_0\,,$$

where $K_D^{dim}$ is the dimerization dissociation constant, $K_D^{mem}$ is the membrane dissociation constant, $c_0$ is a reference concentration (1 molar), $R$ is the gas constant, $T$ is the temperature in Kelvins and $s_0$ is a reference entropy (note that $R$, $T$, $c_0$ and $s_0$ cancel out in the equilibrium condition). Thus, for a given amount of total protein $c_{tot}$, and given values of the dissociation constants $K_D^{dim}$ and $K_D^{mem}$, equilibrium membrane and cytoplasmic concentrations can be calculated according to the equilibrium condition (2) and conservation law (1). Then, within each compartment, monomer and dimer concentrations are given by the following expressions (see supplemental model description for full derivation):

$$c_1 = \frac{K_D^{dim}}{4}\left(\sqrt{1 + \frac{8c}{K_D^{dim}}} - 1\right) \qquad c_2 = \frac{K_D^{dim}}{4}\left(\frac{4c}{K_D^{dim}} - \sqrt{1 + \frac{8c}{K_D^{dim}}} + 1\right)$$

*Scoring cooperativity*

In Figure 3B, cooperativity was calculated by solving systems at equilibrium with $a = 5$ nm, $\psi = 0.174$ µm$^{-1}$ (Goehring et al., 2011b) and $c_{tot}$ varying from 27 nM to 0.27 nM, and performing linear regression on log-transformed equilibrium concentrations:

$$\log_{10}(c_m) = \alpha \log_{10}(c_c) + \beta$$

with the slope ($\alpha$) given as the cooperativity score.

*Fitting model to in vivo PAR-2 data*

To fit this model to our in vivo PAR-2 rundown data, concentrations were first converted from arbitrary units to a best estimate of absolute concentration. To do so, we made use of previous measurements by Gross et al. (2018) which estimate cytoplasmic PAR-2 concentrations in wild type polarized cells to be 10.4 nM, and normalized our concentration measurements accordingly. Wild type and L109R data were then fit simultaneously to a model in which $K_D^{mem}$ is shared between wild type and L109R, with $K_D^{dim}$ (L109R) as a free parameter and $K_D^{dim}$ (wt) either free (Figure 3F) or constrained to the value experimentally determined by AUC (Figure S6). By default we use $a = 5$ nm, however given uncertainty over the true value of $a$, we performed additional fits with $a = 0.5$ nm and 50 nm for comparison (Figure S6, Table S4).

*Six species thermodynamic model*

To include an internal membrane compartment, two additional species were added representing internal membrane bound monomers ($n_1$) and dimers ($n_2$), leading to the new conservation term

$$c_{\text{tot}} = c_c + a(\psi c_m + \phi c_n)$$

where $c_n$ $(= c_{n_1} + c_{n_2})$ is the concentration in the internal membrane compartment, and $\phi$ is the internal membrane surface area to cytoplasmic volume ratio (for simplicity, we assume that $\psi = \phi = 0.087$ µm$^{-1}$ (=0.174/2, reflecting plasma membrane availability in the posterior half)). Equilibrium is given by the new condition:

$$\mu_c = \mu_m = \mu_n$$

with $\mu_n$ given as follows (see supplemental model description for full derivation):

$$\frac{\mu_n}{RT} = \ln\left(\frac{c_n}{c_0}\right) + \ln(K_{\text{D}}^{\text{int}}) - \frac{1}{2}\ln\left(1 + \frac{4c_n}{K_{\text{D}}^{\text{dim}}} + \sqrt{1 + \frac{8c_n}{K_{\text{D}}^{\text{dim}}}}\right) + 1 + s_0\,,$$

where $K_{\text{D}}^{\text{int}}$ is the dissociation constant for internal membranes. For our simulations we used $c_{\text{tot}} = 27$ nM, based on previous estimates of the cytoplasmic PAR-2 concentration in polarized wild type cells (Gross et al., 2018) and our estimate of the cytoplasmic fraction of PAR-2 in these conditions (Table S2).

**Modelling - kinetic model**

We extend our thermodynamic model using transition state theory to derive the following concentration dependent on and off rates (see supplemental model description for full derivation):

$$k_{\text{on}} = \frac{\tilde{\Lambda}}{\sqrt{1 + \frac{4c_c}{K_{\text{D}}^{\text{dim}}} + \sqrt{1 + \frac{8c_c}{K_{\text{D}}^{\text{dim}}}}}} \qquad k_{\text{off,m}} = \frac{\tilde{\Lambda} K_{\text{D}}^{\text{mem}}}{\sqrt{1 + \frac{4c_m}{K_{\text{D}}^{\text{dim}}} + \sqrt{1 + \frac{8c_m}{K_{\text{D}}^{\text{dim}}}}}} \qquad k_{\text{off,n}} = \frac{\tilde{\Lambda} K_{\text{D}}^{\text{int}}}{\sqrt{1 + \frac{4c_n}{K_{\text{D}}^{\text{dim}}} + \sqrt{1 + \frac{8c_n}{K_{\text{D}}^{\text{dim}}}}}}$$

where $c_c$, $c_m$, and $c_n$ are molar concentrations in the cytoplasmic, plasma membrane and internal membrane compartments, and $K_{\text{D}}^{\text{dim}}$, $K_{\text{D}}^{\text{mem}}$, $K_{\text{D}}^{\text{int}}$ are dissociation constants for dimerization, plasma membrane dissociation and internal membrane dissociation as previously defined. $\tilde{\Lambda}$ is a kinetic pre-factor that scales the rates according to kinetic details (see supplemental model description for details). We used experimentally determined off-rate measurements to calibrate $\tilde{\Lambda}$ for PAR-2. FRAP measurements put the plasma membrane unbinding rate in wild type polarized cells at 0.0073 s$^{-1}$ (Goehring et al., 2011a). With $K_{\text{D}}^{\text{dim}} = 358$ µM (AUC), $K_{\text{D}}^{\text{mem}}$ $= 10^{-2.39}$ (model fit), and $c_m = 47.8$ $\mu$M (our quantification of mean plasma membrane concentration in polarized cells assuming $a = 5$ nm), this gives $\tilde{\Lambda} = 43.1$ M s$^{-1}$. Using these concentration-dependent rate expressions, ODE systems were set up with the following governing equations:

$$\frac{dc_c}{dt} = a[\psi(-k_{\text{on}} c_c + k_{\text{off,m}} c_m) + \phi(-k_{\text{on}} c_c + k_{\text{off,n}} c_n)]$$
$$\frac{dc_m}{dt} = k_{\text{on}} c_c - k_{\text{off,m}} c_m$$
$$\frac{dc_n}{dt} = k_{\text{on}} c_c - k_{\text{off,n}} c_n$$

Systems were initiated from an equilibrium state with $K_{\text{D}}^{\text{mem}} = 1$ and $c_{\text{tot}} = 27$ nM. At time zero, $K_{\text{D}}^{\text{mem}}$ was decreased to simulate the onset of posterior plasma membrane availability, and ODE systems were simulated, with $\psi = \phi = 0.087$ µm$^{-1}$, $a = 5$ nm and $\tilde{\Lambda}$ $= 43.1$ M s$^{-1}$.

### Supplemental model description

#### 1 Introduction

In this document, we provide a detailed description of the thermodynamic approaches used to model PAR-2 membrane association and dimerization. We begin by building the equilibrium model described in Figure 3 of the main text, a four species model with two compartments representing the plasma membrane and cytosol. We then extend this model, firstly to consider the system outside of equilibrium, and secondly to add a third compartment representing internal membranes, both of which form the basis of the analysis in Figure 5 of the main text.

#### 2 Equilibrium Model

We aim to describe the equilibrium behavior of PAR-2, taking into account both dimerization and membrane association. To do so, we begin by building separate thermodynamic descriptions for dimerization and membrane association, before describing a full model which contains both features.

##### 2.1 Thermodynamics of dimerization

First, we consider a non-interacting system containing a mixture of monomeric and dimeric proteins and solvent in a single compartment. In order to study the equilibrium condition, we begin by describing the energetic and entropic contributions, from which we will derive the free energy of the system. From the free energy, we will then derive chemical potentials associated with monomeric and dimeric protein. Then, equating such chemical potentials will provide the dimerization equilibrium condition.

###### 2.1.1 Energy

Consider a system composed of a mixture of monomeric and dimeric proteins and a solvent. Since proteins convert between a monomeric and a dimeric state, the total number concentration of protein is conserved. Thus the following conservation law holds

$$C_1 + C_2 = C_{\text{tot}} \,, \tag{1}$$

where $C_1$, $C_2$ are the number concentrations of proteins in the monomer and dimer state, respectively, and $C_{tot}$ is the total number concentration of proteins. (Note that $C_2$ corresponds to the number of proteins in the dimeric state, not the number of dimers). Since we are considering the system to be non interacting, the energy density of a configuration can be calculated as

$$\frac{H}{V} = RT(c_1\omega_1 + c_2\omega_2) \,, \tag{2}$$

where $\omega_1$, $\omega_2$ are dimensionless internal energies of protein in the monomer and dimer states respectively, $V$ is the total volume of the system (including solvent), $c_i = C_i/N_A$ are the molar concentrations and $R = k_\text{B} N_\text{A}$ is the gas constant.

###### 2.1.2 Entropy

Starting from the result of Flory [1], we can write the entropy density in terms of number concentrations of protein in monomer and dimer states as follows

$$\Delta s = -k_\text{B}\left(C_1 \ln(C_1\nu) + \frac{C_2}{2}\ln(C_2\nu) + (\frac{1}{\nu} - C_1 - C_2)\ln\left(1 - \nu C_1 - \nu C_2\right)\right) \,, \tag{3}$$

which depends on the volume of a protein $\nu$. Considering the system to be in the *dilute limit* $\nu(C_1 + C_2) \ll 1$

$$\Delta s = -k_\mathrm{B}\left( C_1 \ln(C_1\nu) + \frac{C_2}{2}\ln(C_2\nu)\right) . \tag{4}$$

Recasting in terms of molar concentrations

$$\Delta s = -R\left( c_1 \ln(c_1 N_\mathrm{A}\nu) + \frac{c_2}{2}\ln(c_2 N_\mathrm{A}\nu)\right) , \tag{5}$$

which we can rewrite as

$$\Delta s = -R\left( c_1 \ln\left(\frac{c_1}{c_0}\right) + \frac{c_2}{2}\ln\left(\frac{c_2}{c_0}\right) + s_0\left(c_1 + \frac{c_2}{2}\right)\right) , \tag{6}$$

where $c_0$ is a reference molar concentration introduced to obtain a more familiar form, and $s_0 = \ln(c_0 N_\mathrm{A}\nu)$ is a reference entropy introduced by the reference molar concentration.

###### 2.1.3 Free energy and chemical potentials

We obtain the free energy density $f = H/V - T\Delta s$ in terms of molar concentrations of protein in monomer and dimer state, still considering the system to be in the dilute limit

$$\frac{f}{RT} = c_1 \ln\left(\frac{c_1}{c_0}\right) + \frac{c_2}{2}\ln\left(\frac{c_2}{c_0}\right) + c_1(\omega_1 + s_0) + c_2\left(\omega_2 + \frac{s_0}{2}\right) . \tag{7}$$

The associated chemical potentials are then obtained by taking the derivative of the free energy with respect to $c_1$ and $c_2$ as follows

$$\mu_1 = \frac{\partial f}{\partial c_1} = RT\left( \ln\left(\frac{c_1}{c_0}\right) + \omega_1 + 1 + s_0\right) \tag{8a}$$

$$\mu_2 = \frac{\partial f}{\partial c_2} = RT\left( \frac{1}{2}\ln\left(\frac{c_2}{c_0}\right) + \omega_2 + \frac{1+s_0}{2}\right) , \tag{8b}$$

where the chemical potentials shown here are chemical potentials associated with one mole of constituents.

###### 2.1.4 Dimerization equilibrium

Considering the dimerization process to be fast, one can assume that dimerization equilibrium is satisfied, i.e.

$$\mu_1 = \mu_2 . \tag{9}$$

Imposing this condition together with the mass conservation condition (1) expressed in terms of molar concentrations

$$c_1 + c_2 = c , \tag{10}$$

we can express the molar concentrations of protein in the monomer and dimer states as functions of the total molar concentration of proteins. We obtain

$$c_1 = \frac{K_\mathrm{D}^\mathrm{dim}}{4}\left(\sqrt{1 + \frac{8c}{K_\mathrm{D}^\mathrm{dim}}} - 1\right) \qquad c_2 = \frac{K_\mathrm{D}^\mathrm{dim}}{4}\left(\frac{4c}{K_\mathrm{D}^\mathrm{dim}} - \sqrt{1 + \frac{8c}{K_\mathrm{D}^\mathrm{dim}}} + 1\right) , \qquad \frac{2c_1^2}{c_2} = K_\mathrm{D}^\mathrm{dim} , \tag{11}$$

upon defining the dimer dissociation constant $K_\mathrm{D}^\mathrm{dim} = 2c_0 e^{2(\omega_2 - \omega_1) - 1 - s_0}$. Then, inserting these expressions into the free energy expression (7), one obtains an effective free energy depending only on the total molar concentration. Taking the derivative of this free energy with respect to $c$, the following non dimensionalized effective chemical potential is obtained

$$\frac{\mu}{RT} = \ln\left(\frac{c}{c_0}\right) + (1 + \omega_1) - \frac{1}{2}\ln\left(1 + \frac{4c}{K_\mathrm{D}^\mathrm{dim}} + \sqrt{1 + \frac{8c}{K_\mathrm{D}^\mathrm{dim}}}\right) + s_0 . \tag{12}$$

#### 2.2 Thermodynamics of membrane association

We now separately describe the thermodynamics of a protein exchanging between the cytosol and membrane. We describe the system as two compartments: a three-dimensional bulk and a membrane which we describe as three dimensional and of small thickness $a$, where $a$ is the microscopic length scale of proteins. As protein exchanges between the two compartments, the total concentration is conserved according to the following conservation law

$$c_c + a\psi c_m = c_{\text{tot}} \, , \tag{13}$$

where $c_m$, $c_c$ are the membrane and cytosol molar concentrations, $\psi = A/V$ is the ratio of membrane surface area over bulk volume, and $c_{\text{tot}}$ is the concentration when all the proteins are in the cytosol. To study the equilibrium we proceed as in the previous section and calculate the membrane and bulk chemical potentials. The entropic contribution will now have the same form in the two compartments, so we obtain the following non dimensionalized ideal gas forms

$$\frac{\mu_m}{RT} = \ln\left(\frac{c_m}{c_0}\right) + \omega_m \quad \frac{\mu_c}{RT} = \ln\left(\frac{c_c}{c_0}\right) + \omega_c \, , \tag{14}$$

neglecting the reference entropies that are just additive constants, and introducing the non-dimensional internal energies $\omega_m$,$\omega_c$. The equilibrium condition is

$$\mu_m = \mu_c \tag{15}$$

implying a constant ratio of protein in the two compartments at equilibrium according to the difference between the two internal energies

$$\frac{c_m}{c_c} = e^{\omega_c - \omega_m} \tag{16}$$

#### 2.3 Full model

Finally, we describe a full model consisting of both dimerization and membrane exchange. In this model there are four protein states: monomer or dimer, membrane bound or in the bulk. To characterize the four states we *assign the internal energies* as follows

- When a protein is bound to the membrane an energy $\omega_m$ is assigned
- When a protein is in dimer state the energy $\omega_d$ is assigned

We assume that dimerization and membrane association are independent, i.e. dimerization energy is the same on the membrane and in the cytosol, and the membrane association energy per protein is the same whether the protein is monomeric or dimeric. Writing explicitly the nondimensional internal energies of the four protein species we obtain

$$\omega_{c,1} = 0 \quad \omega_{c,2} = -\omega_d \quad \omega_{m,1} = -\omega_m \quad \omega_{m,2} = -\omega_d - \omega_m \, . \tag{17}$$

Then, imposing dimerization equilibrium in each compartment, we insert the membrane and bulk internal energies into the general effective chemical potential (12), obtaining an *effective two state model* described by the following non dimensionalized membrane and bulk chemical potentials

$$\frac{\mu_c}{RT} = \ln\left(\frac{c_c}{c_0}\right) - \frac{1}{2}\ln\left(1 + \frac{4c_c}{K_{\text{D}}^{\text{dim}}} + \sqrt{1 + \frac{8c_c}{K_{\text{D}}^{\text{dim}}}}\right) + 1 + s_0 \tag{18a}$$

$$\frac{\mu_m}{RT} = \ln\left(\frac{c_m}{c_0}\right) + \ln(K_{\text{D}}^{\text{mem}}) - \frac{1}{2}\ln\left(1 + \frac{4c_m}{K_{\text{D}}^{\text{dim}}} + \sqrt{1 + \frac{8c_m}{K_{\text{D}}^{\text{dim}}}}\right) + 1 + s_0 \, , \tag{18b}$$

where $K_{\text{D}}^{\text{dim}} = 2c_0 e^{-2\omega_d - 1 - s_0}$ and $K_{\text{D}}^{\text{mem}} = e^{-\omega_m}$. For a given total amount of protein $c_{tot}$, and given values of the dissociation constants $K_{\text{D}}^{\text{dim}}$ and $K_{\text{D}}^{\text{mem}}$, equilibrium membrane and cytosolic concentrations can be calculated according to the equilibrium condition $\mu_m = \mu_c$ and the conservation law (13). (Note that at equilibrium the additive term $1 + s_0$ cancels out so can be neglected). Then, once overall membrane and cytosol concentrations are calculated, concentrations of monomer and dimer within each compartment can be calculated according to the relationship shown previously (11).

### 3 Kinetic model

To explicitly evaluate membrane exchange rates we study the effective two state model in a non-equilibrium situation. We can imagine bringing this system out of equilibrium, for example placing all the system components in one of the two states, and letting it evolve. As the system relaxes towards equilibrium detailed balance holds [3]

$$\frac{s_{\text{on}}}{s_{\text{off}}} = \exp\left(\frac{\mu_c - \mu_m}{RT}\right), \tag{19}$$

where we define the individual attachment and detachment fluxes $s_{\text{on}}$, $s_{\text{off}}$, that together define the total flux $s = s_{\text{on}} - s_{\text{off}}$. To evaluate the individual fluxes we can use transition state theory [2]. We can think of a free energy landscape composed of two local minima, corresponding to the membrane and bulk state, separated by an energy barrier. A simple approach is, upon assuming that the metastable states are in a local equilibrium condition and considering the flux small and constant such that it can be considered stationary, to estimate flux using a Smoluchowski equation. This approach leads to the result that the rate with which components can overcome the energy barrier is proportional to the Bolztmann factor associated with the energy barrier

$$s_{\text{on}} = k \exp\left[-\frac{(\mu_{\text{max}} - \mu_c)}{RT}\right] = \Lambda \exp\left(\frac{\mu_c}{RT}\right) \tag{20a}$$

$$s_{\text{off}} = k \exp\left[-\frac{(\mu_{\text{max}} - \mu_m)}{RT}\right] = \Lambda \exp\left(\frac{\mu_m}{RT}\right), \tag{20b}$$

where $k$ is a factor related to the kinetic details of the system. We know this to be equal for the two metastable states due to the detailed balance condition (this is not true in general since $k$ depends form the details of the local energy minimum). Together with the Boltzmann factor associated with the local energy maxima, this defines the prefactor $\Lambda$. Inserting our expressions for the chemical potentials Eq. (18), we obtain

$$s_{\text{on}} = \frac{\tilde{\Lambda} c_c}{\sqrt{1 + \frac{4c_c}{K_{\text{D}}^{\text{dim}}} + \sqrt{1 + \frac{8c_c}{K_{\text{D}}^{\text{dim}}}}}} \qquad s_{\text{off}} = \frac{\tilde{\Lambda} K_{\text{D}}^{\text{mem}} c_m}{\sqrt{1 + \frac{4c_m}{K_{\text{D}}^{\text{dim}}} + \sqrt{1 + \frac{8c_m}{K_{\text{D}}^{\text{dim}}}}}} \tag{21}$$

where $\tilde{\Lambda} = \Lambda e^{s_0+1}/c_0$. Finally, assuming that these fluxes have a mass action kinetics inspired form $s = k_{\text{on}} c_c - k_{\text{off}} c_m$, we find the concentration dependent rates

$$k_{\text{on}} = \frac{\tilde{\Lambda}}{\sqrt{1 + \frac{4c_c}{K_{\text{D}}^{\text{dim}}} + \sqrt{1 + \frac{8c_c}{K_{\text{D}}^{\text{dim}}}}}} \qquad k_{\text{off}} = \frac{\tilde{\Lambda} K_{\text{D}}^{\text{mem}}}{\sqrt{1 + \frac{4c_m}{K_{\text{D}}^{\text{dim}}} + \sqrt{1 + \frac{8c_m}{K_{\text{D}}^{\text{dim}}}}}}. \tag{22}$$

### 4 Model incorporating internal membranes

We also explore a model in which proteins have access to a third compartment representing internal membranes. Introducing this compartment leads to the new conservation term

$$c_c + a(\psi c_m + \phi c_n) = c_{\text{tot}}, \tag{23}$$

where $c_n$ is the concentration in the internal membrane compartment and $\phi$ ratio of internal membrane surface area over bulk volume. As before, we consider the mixture of monomeric and dimeric proteins on internal membranes to be in dimerization equilibrium, obtaining the following chemical potential

$$\frac{\mu_n}{RT} = \ln\left(\frac{c_n}{c_0}\right) + \ln(K_{\text{D}}^{\text{int}}) - \frac{1}{2}\ln\left(1 + \frac{4c_n}{K_{\text{D}}^{\text{dim}}} + \sqrt{1 + \frac{8c_n}{K_{\text{D}}^{\text{dim}}}}\right) + 1 + s_0, \tag{24}$$

with the internal membrane dissociation constant $K_{\text{D}}^{\text{int}}$. For a given total amount of protein $c_{tot}$, and given values of the dissociation constants $K_{\text{D}}^{\text{dim}}$, $K_{\text{D}}^{\text{mem}}$ and $K_{\text{D}}^{\text{int}}$, equilibrium concentrations in each of the three compartments are calculated according to the mass conservation term and the new equilibrium condition

$$\mu_c = \mu_m = \mu_n. \tag{25}$$

Evaluating the out of equilibrium exchange rates between the cytosol and internal membranes, we obtain the rate of detachment from the internal membranes as

$$k_{\text{off,n}} = \frac{\tilde{\Lambda} K_{\text{D}}^{\text{int}}}{\sqrt{1 + \frac{4c_n}{K_{\text{D}}^{\text{dim}}} + \sqrt{1 + \frac{8c_n}{K_{\text{D}}^{\text{dim}}}}}}. \tag{26}$$
